# Mutational load, inbreeding depression and heterosis in subdivided populations

**DOI:** 10.1101/352146

**Authors:** Brian Charlesworth

## Abstract

This paper examines the extent to which empirical estimates of inbreeding depression and inter-population heterosis in subdivided populations, as well as the effects of local population size on mean fitness, can be explained in terms of estimates of mutation rates, and the distribution of selection coefficients against deleterious mutations provided by population genomics data. Using results from population genetics models, numerical predictions of the genetic load, inbreeding depression and heterosis were obtained for a broad range of selection coefficients and mutation rates. The models allowed for the possibility of very high mutation rates per nucleotide site, as is sometimes observed for epiallelic mutations. There was fairly good quantitative agreement between the theoretical predictions and empirical estimates of heterosis and the effects of population size on genetic load, on the assumption that the deleterious mutation rate per individual per generation is approximately one, but there was less good agreement for inbreeding depression. Weak selection, of the order of magnitude suggested by population genomic analyses, is required to explain the observed patterns. Possible caveats concerning the applicability of the models are discussed.

## 1 INTRODUCTION

There is now a substantial body of data on mean population fitness, inbreeding depression and heterosis in subdivided populations, reviewed by Byers & Waller, (1999), Keller & Waller (2002) and Leimu, Mutikainen, Koricheva *&* Fischer (2006). A number of relevant theoretical investigations have also been carried out (Escobar, Nicot & David, 2008; García-Dorado 2007, 2008; Glémin, Ronfort & Bataillon, 2003; Roze, 2015; Roze & Rousset, 2004; Theodorou & Couvet, 2002; Whitlock, Ingvarsson & Hatfield, 2000). These have shown that the mutational load and inbreeding depression are generally increased by high migration rates and large local population sizes, whereas heterosis in between-population crosses is reduced. While the theoretical expectations are in qualitative agreement with the empirical results, no attempt has been made to determine whether the data quantitatively match theoretical predictions based on estimates of mutation rates and the distribution of fitness effects of new deleterious mutations obtained from genomic data, which have been reviewed by Keightley (2012) and Charlesworth (2015).

Furthermore, the theoretical work has assumed that mutation rates per locus or nucleotide site are small, relative to the strength of selection against deleterious alleles. In addition, the forward mutation rate (from wild-type to mutant) is usually assumed to be much greater than the backward mutation rate (from mutant to wild-type), the latter often being ignored. The first assumption is usually well-justified for mutations affecting protein sequences and strongly constrained functional non-coding sequences. The second assumption applies to the totality of mutations that affect a functional unit such as a coding sequence, provided that selection is sufficiently strong in relation to drift that mutant alleles are close to deterministic equilibrium. This is because there are many ways in which a wild-type sequence can mutate to a sequence with impaired function, but the reversion of a mutated sequence requires a change at the site of the original mutation. However, if we are considering individual nucleotide sites under weak selection, models that include drift and reverse mutation need to be used (Charlesworth, 2013; Kimura, Maruyama & Crow, 1963; Kondrashov, 1995).

The importance of examining the consequences of relaxing these two assumptions is brought out by recent studies of variation in epigenetic marks in plants such as *Arabidopsis thaliana*, especially methylation at CG sites. These have found rates of gain and loss of marks at individual nucleotide sites that are about 5,000 times higher than the DNA sequence mutation rate, with a strong bias towards loss of methylation (Quadrana & Colot, 2016; Van De Graaf *et al.*, 2015; Vidalis, Zivkovic, Wardenaar, Roquis, Tellier & Johannes, 2016). If fitness is affected by epiallelic variants at numerous sites in the genome, the above assumptions are likely to be violated (Charlesworth & Jain, 2014).

In this paper, theoretical expectations are derived for the mutational load, inbreeding depression and between-population heterosis in subdivided populations, allowing for the possibility of some level of inbreeding, as well as for potentially very high mutation rates per nucleotide site, using an extension of the method described by Charlesworth & Jain (2014). The results are related to data on the relevant parameters obtained from the literature, using parameters of mutation and selection derived from genomic studies. It is shown that there is fairly good quantitative agreement between the theoretical predictions and estimates of heterosis and the effects of local population size on genetic load, but less good agreement for inbreeding depression. The ground is prepared by describing the basic model of a single randomly mating population.

## 2. THEORETICAL METHODS AND RESULTS

### 2.1 Basic model of a randomly mating population of infinite size

The standard population genetic model of selection and mutation at a biallelic autosomal locus is assumed (Charlesworth & Charlesworth, 2010, Chap. 4), The relative fitnesses of the genotypes A_1_A_1_, A_1_A_2_ and A_2_A_2_ are 1, 1 − *hs* and 1 − *s*; the frequencies of A_1_ and A_2_ in a given generation are *p* and *q*, respectively. The forward and backward mutation rates (A_1_ to A_2_ and A_2_ to A_1_, respectively) are *u* and *v*. It is convenenient to write *u* = *κv,* where *κ* measures the extent of mutational bias. If *κ* > 1, mutation is biased towards the production of deleterious alleles; if *κ* < 1, the bias is toward wild-type alleles.

Ignoring second-order terms in *s*, *u* and *v*, the change in *q* per generation in a randomly mating, discrete generation population is given by:

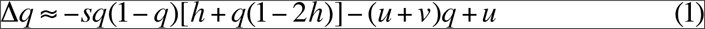

The mean fitness of the population is:

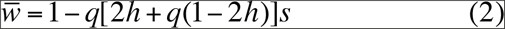

and the genetic load is:

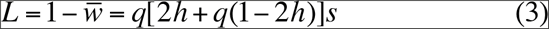

The equilibrium frequency of A_2_, *q*\*, is found by equating equation (1) to zero. Further details are described in the Supplementary Material, section 1. The corresponding equilibrium load, *L*\*, given by this procedure is always smaller than the low mutation rate approximation, 2*u*, for non-recessive autosomal mutations (Haldane, 1937), but approaches 2*u* when *hs* ≫ (*u* + *v*).

Table S1 and Figure S1 show some numerical examples of the dependence of *q*\* and *L*\* on the selection and mutation parameters, which are discussed in the Supplementary Material, section 1. In the next section, the effects of finite population size are examined.

### 2.2 Stochastic results for a randomly mating population

Here, the load is measured by taking its expected value over the distribution of allele frequencies, following the approach used by Kimura *et al.* (1963) and Bataillon & Kirkpatrick (2000) for the low mutation rate case. Following Kimura *et al.* (1963), insertion of equation (2) into the general expression for the stationary probability distribution of allele frequencies at a single locus (Wright, 1937a; Charlesworth & Charlesworth, 2010, Chap. 5) yields an expression for the probability density of *q* in a randomly mating population with effective population size *N*_*e*_:

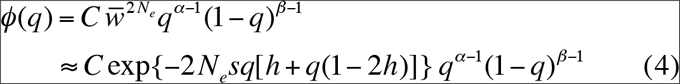

where *N*_*e*_ is the effective population size, *α* = 4*N*_*e*_*u*, *β* = 4*N*_*e*_*v*, and *C* is a constant of integration that ensures that the integral of *ϕ*(*q*) between 0 and 1 is unity.

Equations (S4) – (S7) of section 2 of the Supplementary Material, show how this expression can be used to obtain explicit formulae for the moments about zero of the allele frequency at mutation-selection-drift equilibrium, using infinite series representations of the relevant integrals. This avoids the need to use either numerical integration or the linearized approximation for the effect of selection, given by equations (S10) and (S11) in the Supplementary Material, section 3. Both of these approaches have been employed in past treatments, e.g. Bataillon & Kirkpatrick (2000); Glémin *et al.* (2003), and Roze & Rousset (2004). For an alternative approach, see García-Dorado (2007, 2008).

Some results for the dependence of the expected load (equation S8) on the selection and mutation parameters are shown in Figure S2. The general picture is very similar to that described by Kimura *et al.* (1963). There is a critical value of the scaled selection parameter, *γ* = 2*N*_*e*_*s*, below which drift and mutation start to overcome selection, leading to a load that is much greater than the deterministic equilibrium value. While high mutation rates tend to reduce the expected load relative to its deterministic value, its absolute value always increases with the mutation rate. As shown below, the same pattern is observed in subdivided populations.

### 2.3 Analytical results for a subdivided population with random mating within local populations

It is useful to consider first some general results on subdivided populations, which do not assume a specific population structure. While similar results have been obtained before (Escobar *et al.*, 2008; Glémin *et al.*, 2003; Roze, 2015; Whitlock *et al.*, 2000), the simple but general derivation given below does not seem to have been published. Here, inbreeding depression is defined as the difference between the expected mean fitness over all demes and the expected fitness of completely homozygous individuals formed from alleles sampled within a deme. Because the effect of a single locus is very small, this measure is equivalent to the *B* coefficient of Morton, Crow & Muller (1956). This coefficient is equal to the regression of the natural logarithm of fitness on the inbreeding coefficient of an individual, and is widely used to describe empirical estimates of the effects of inbreeding, e.g. Charlesworth & Charlesworth (1987) and Charlesworth & Charlesworth (2010, Chap. 4). On the assumption of multiplicative fitness effects of multiple independent loci, *B* is given by the sum of the individual locus values.

For a single locus, whose effect on mean fitness can be assumed to be very small, the expected load suffered by a deme (equation S8) can be written as:

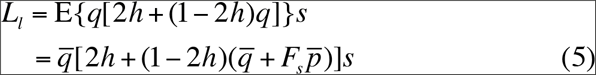

where *F*_*s*_ is the inbreeding coefficient that measures population differentiation at a locus subject to mutation and selection, overbars indicate expectations over demes, and the subscript *l* indicates that *L*_*l*_ is the value for a single locus or nucleotide site.

This equation brings out the important fact that the mean load is affected by both the effects of drift on the mean of *q* over all demes, and on the extent of variation in *q* among demes when *h* ≠ ½.

The corresponding measure of the expected inbreeding depression is:

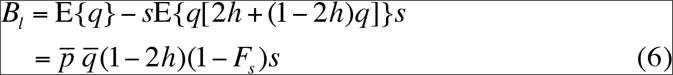

This shows that population subdivision tends to reduce the mean inbreeding depression when *h* < ½, even if the mean frequencies of mutant alleles are unchanged from the random mating case.

The value of *B*_*l*_ under classical mutation-selection equilibrium with *u* ≪ *hs* is 2*u*[1/(2*h*) − 1], provided that *h* > 0 (Charlesworth & Charlesworth, 1987). It is helpful to use the quantity *B*_*rel*_ = *B*_*l*_ /2*u* as an estimate of the value of the sum of the *B*_*l*_ over loci relative to the diploid genome-wide deleterious mutation rate, *U*, which is equal to the sum of the values of 2*u* over all loci. A relative mean load, *L*_*rel*_ = *L*_*l*_/(2*u*), can be defined in the same way.

Similarly, the expected between-deme heterosis (*H*_*l*_) caused by a single locus is given by the expected difference between the fitnesses of random matings among individuals from different demes and random matings within demes (Glémin *et al.*, 2003). This is given by:

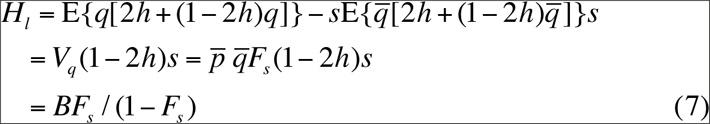

where *V*_*q*_ is the variance among demes in allele frequency at the selected locus. Again, use of *H*_*rel*_ = *H*_*l*_ /(2*u*) allows the sum of *H*_*l*_ values over loci to be compared with *U*.

These expressions bring out clearly that the extent of population subdivision, as measured by *F*_*s*_, has complementary effects on *B*_*l*_ and *H*_*l*_. *F*_*s*_ is expected to be smaller than the genealogical measure of subdivision, *F*_*ST*_, which assumes neutral transmission (Glémin *et al.*, 2003). However, since *F*_*ST*_ is relatively easy to measure using genetic markers, it can be used as a rough guide as what to expect from equations (6) and (7) (Charlesworth & Charlesworth, 2010, pp.361-362). The numerical examples described below for the infinite island model suggest that the neutral approximation works well when *s* ≪ *m*, as is predicted by equation (S12).

Further results can most be easily obtained using Wright’s island model of population structure with an infinite number of demes (Wright, 1937b), which assumes that the metapopulation is composed of a large number of randomly mating demes, each with effective size *N*_*e*_. Each deme receives a constant fraction *m* of its gene pool from the population as a whole, with an expected frequency *q*_*t*_ of allele A_2_ among the migrants that is equal to the mean allele frequency for the whole metapopulation, which is assumed to be fixed. This corresponds to soft selection in the sense of (Whitlock *et al.*, 2000).

The stationary distribution of allele frequencies among demes under the infinite island model (Wright, 1937b) can be found from the analogue of equation (4):

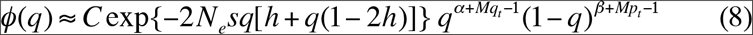

where *M* is the scaled migration parameter, 4*N*_*e*_*m*, and *p*_*t*_= 1 − *q*_*t*_.

However, in general *q*_*t*_ cannot be equated to the deterministic equilibrium value, *q*\*. It can be found by iteratively determining the value of *q*_*t*_ that corresponds to the mean of *q* over the distribution given by equation (8), using a simple extension of the method described in the Supplementary Material, section 2. Linearized approximations are provided by the equivalents of equations (S10) and (S11).

### 2.4 Numerical results for a subdivided population with random mating within local populations

The results described here assume weak selection, with *s* = 2.5 × 10^−3^. There are two reasons for this; first, unless deme sizes are extremely small (≤ 100), the effects of drift will be minor when selection coefficients are an order of magnitude or more greater than this (Kimura *et al.*, 1963). Second, recent population genomic studies of *Drosophila* suggest that there is a wide distribution of selection coefficients against deleterious nonsynonymous mutations and mutations in functional non-coding sequences, with a mean selection coefficient (*hs*) of the order of 10^−3^ for nonsynonymous mutations, and a much smaller value for mutations in functional non-coding sequences (Campos, Zhao & Charlesworth, 2017; Kousathanas & Keightley, 2013). The outcrossing flowering plant, *Capsella grandiflora*, has a similar estimated distribution of the fitness effects of nonsynonymous mutations to that in *Drosophila* (Slotte, Foxe, Hazzouri & Wright, 2010). With the highly left-skewed distributions suggested by these analyses, most deleterious mutations will have considerably smaller *s* values than the mean. To be conservative in the sense of underestimating the effects of drift, *s* = 2.5 × 10^−3^ was chosen for the numerical analyses.

Table 1 shows some examples of the effects of varying the deme size, mutation rate and scaled migration rate on the exact and approximate values of the expectations of the parameters of interest. As described in section 2.3, these are divided by the diploid mutation rate, 2*u*, as indicated by the subscript *rel*. There are several points of interest. First, for the smaller deme size the exact expected load is considerably higher than 2*u* for the lower pair of values of the scaled migration rate *M,* reflecting the effect of drift in causing deleterious alleles to drift to high frequencies when 2*N*_*e*_*s* is of order 1 or less. This effect is not seen with the linear approximation, because this assumes that the mean of *q* is equal to the deterministic equilibrium value, *q*\*.

**Table 1.**
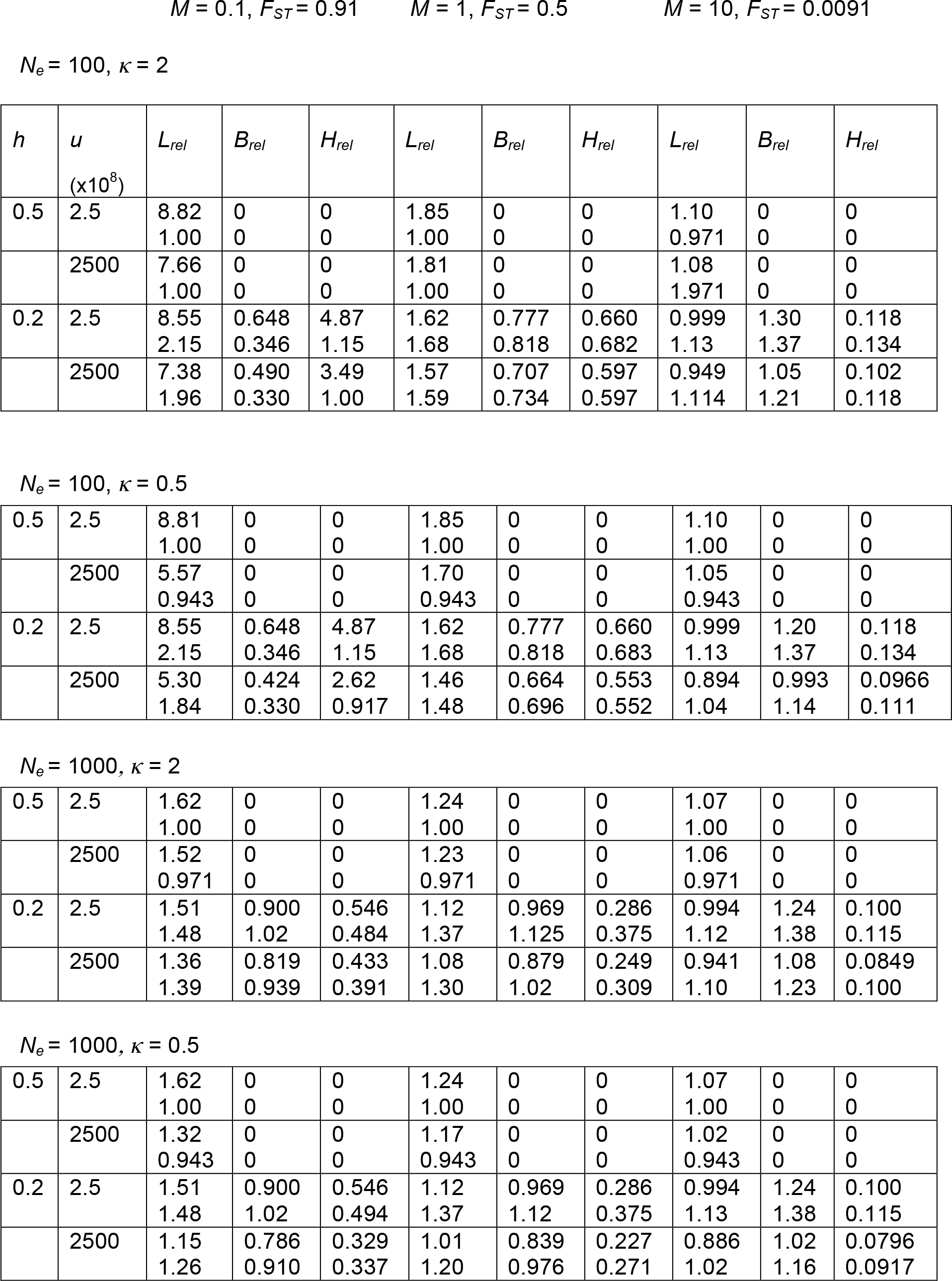
Genetic loads, inbreeding depression and heterosis in a subdivided population. All results are expressed relative to the diploid mutation rate, *2u*, as indicated by the subscript *rel*. The expected loads are denoted by *L*_*rel*_. The inbreeding depression values (*B*_*rel*_) were calculated from equation (6), and the heterosis values (*H*_*rel*_) from equation (7). The selection coefficient *s* was 2.5 × 10^−3^. The upper items in the cells are the exact expectations over the probability distribution of local allele frequencies, using equations (S4) - (S6), where *α* and *β* in the latter are replaced by *α* + *Mq*_*t*_ and *β* + *Mp*_*t*_, respectively. The lower items in the cells were calculated from the linearized approximation to this distribution, using equations (S11) and (S12).

| $h$ | $u$<br>($\times 10^8$) | $L_{rel}$ | $B_{rel}$ | $H_{rel}$ | $L_{rel}$ | $B_{rel}$ | $H_{rel}$ | $L_{rel}$ | $B_{rel}$ | $H_{rel}$ |
| --- | --- | --- | --- | --- | --- | --- | --- | --- | --- | --- |
| 0.5 | 2.5 | 8.82<br>1.00 | 0<br>0 | 0<br>0 | 1.85<br>1.00 | 0<br>0 | 0<br>0 | 1.10<br>0.971 | 0<br>0 | 0<br>0 |
|  | 2500 | 7.66<br>1.00 | 0<br>0 | 0<br>0 | 1.81<br>1.00 | 0<br>0 | 0<br>0 | 1.08<br>1.971 | 0<br>0 | 0<br>0 |
| 0.2 | 2.5 | 8.55<br>2.15 | 0.648<br>0.346 | 4.87<br>1.15 | 1.62<br>1.68 | 0.777<br>0.818 | 0.660<br>0.682 | 0.999<br>1.13 | 1.30<br>1.37 | 0.118<br>0.134 |
|  | 2500 | 7.38<br>1.96 | 0.490<br>0.330 | 3.49<br>1.00 | 1.57<br>1.59 | 0.707<br>0.734 | 0.597<br>0.597 | 0.949<br>1.114 | 1.05<br>1.21 | 0.102<br>0.118 |
| $$N_e = 100, \kappa = 0.5$$ |  |  |  |  |  |  |  |  |  |  |
| 0.5 | 2.5 | 8.81<br>1.00 | 0<br>0 | 0<br>0 | 1.85<br>1.00 | 0<br>0 | 0<br>0 | 1.10<br>1.00 | 0<br>0 | 0<br>0 |
|  | 2500 | 5.57<br>0.943 | 0<br>0 | 0<br>0 | 1.70<br>0.943 | 0<br>0 | 0<br>0 | 1.05<br>0.943 | 0<br>0 | 0<br>0 |
| 0.2 | 2.5 | 8.55<br>2.15 | 0.648<br>0.346 | 4.87<br>1.15 | 1.62<br>1.68 | 0.777<br>0.818 | 0.660<br>0.683 | 0.999<br>1.13 | 1.20<br>1.37 | 0.118<br>0.134 |
|  | 2500 | 5.30<br>1.84 | 0.424<br>0.330 | 2.62<br>0.917 | 1.46<br>1.48 | 0.664<br>0.696 | 0.553<br>0.552 | 0.894<br>1.04 | 0.993<br>1.14 | 0.0966<br>0.111 |
| $$N_e = 1000, \kappa = 2$$ |  |  |  |  |  |  |  |  |  |  |
| 0.5 | 2.5 | 1.62<br>1.00 | 0<br>0 | 0<br>0 | 1.24<br>1.00 | 0<br>0 | 0<br>0 | 1.07<br>1.00 | 0<br>0 | 0<br>0 |
|  | 2500 | 1.52<br>0.971 | 0<br>0 | 0<br>0 | 1.23<br>0.971 | 0<br>0 | 0<br>0 | 1.06<br>0.971 | 0<br>0 | 0<br>0 |
| 0.2 | 2.5 | 1.51<br>1.48 | 0.900<br>1.02 | 0.546<br>0.484 | 1.12<br>1.37 | 0.969<br>1.125 | 0.286<br>0.375 | 0.994<br>1.12 | 1.24<br>1.38 | 0.100<br>0.115 |
|  | 2500 | 1.36<br>1.39 | 0.819<br>0.939 | 0.433<br>0.391 | 1.08<br>1.30 | 0.879<br>1.02 | 0.249<br>0.309 | 0.941<br>1.10 | 1.08<br>1.23 | 0.0849<br>0.100 |
| $$N_e = 1000, \kappa = 0.5$$ |  |  |  |  |  |  |  |  |  |  |
| 0.5 | 2.5 | 1.62<br>1.00 | 0<br>0 | 0<br>0 | 1.24<br>1.00 | 0<br>0 | 0<br>0 | 1.07<br>1.00 | 0<br>0 | 0<br>0 |
|  | 2500 | 1.32<br>0.943 | 0<br>0 | 0<br>0 | 1.17<br>0.943 | 0<br>0 | 0<br>0 | 1.02<br>0.943 | 0<br>0 | 0<br>0 |
| 0.2 | 2.5 | 1.51<br>1.48 | 0.900<br>1.02 | 0.546<br>0.494 | 1.12<br>1.37 | 0.969<br>1.12 | 0.286<br>0.375 | 0.994<br>1.13 | 1.24<br>1.38 | 0.100<br>0.115 |
|  | 2500 | 1.15<br>1.26 | 0.786<br>0.910 | 0.329<br>0.337 | 1.01<br>1.20 | 0.839<br>0.976 | 0.227<br>0.271 | 0.886<br>1.02 | 1.02<br>1.16 | 0.0796<br>0.0917 |

With *M* = 10, the load approaches the classical deterministic value, 2*u*, and is even slightly lower than this when *h* = 0.2, reflecting a weak effect of purging of partially recessive alleles from large but finite populations (Wang, Hill, Charlesworth & Charlesworth, 1999). As is also the case for a single population (sections 2.1 and 2.2), the higher mutation rate is associated with smaller values of the load relative to 2*u,* especially with mutational bias towards favoured alleles, but the absolute mean load is much higher than with the lower mutation rate, even with *N*_*e*_ = 100 and *M* = 0.1. When *h* = 0.2, the linearized approximation of equation (S12) gives quite a good fit to *L*_*rel*_, *B*_*rel*_ and *H*_*rel*_ for the cases with *M* ≥ 1 and *N*_*e*_ = 100; the fit is good for all three values of *M* when *N*_*e*_ = 1000, reflecting the relatively small effect of drift in this case. For *M* = 0.1, there are substantial discrepancies between the approximate and exact values, reflecting the large departures from *q*\*.

Second, as expected from equations (6) and (7), the values of *B*_*rel*_ and *H*_*rel*_ for *h* = 0.2 respond in opposite directions to changes in *N*_*e*_ and *M*, with the level of heterosis being similar to, or even greater than, 2*u* when deme size is small and *M* ≤ 1, but is much less than 2*u* when deme size when deme size is large or *M ≥* 10. *B*_*rel*_ for *h* = 0.2 is mostly well below its deterministic value of 1.5 for the smaller pair of values of *M*, but approaches this value when *M* = 10. *F*_*s*_ was found to be close to the genealogical value, *F*_*ST*_ = 1/(1 + *M*), for most of the examples in Table 1, except for *N*_*e*_ = 1000 and *M* < 10 as well as 2*N*_*e*_*s* ≫ *M*.

There could thus be a substantial expected level of heterosis in interpopulation crosses, even with moderately large *N*_*e*_ or *M*, when selection is of the strength considered here. With *U* = 1, the sum of the *H* values over all sites with the selection and mutation parameters in Table 1 would be approximately 0.60 with *h* = 0.2, *N*_*e*_ =100 and *M* = 1, and approximately 0.10 even with *N*_*e*_ =1000 and *M* = 10.

Finally, the high mutation rate cases in Table 1 all give values of *L*_*rel*_, *B*_*rel*_ and *H*_*rel*_ that are smaller than the corresponding values with low mutation rates; the differences are more marked for *L*_*rel*_ than the other variables. However, the absolute values of all three variables are always much greater with the high mutation rate. Highly mutable, non-neutral epimutations could, therefore, cause high levels of mutational load, inbreeding depression and heterosis.

### 2.5 Analytical and numerical results for subdivided populations with inbreeding

As discussed in section 3 below, much of the evidence for inbreeding depression and interpopulation heterosis in subdivided populations comes from populations with a mixture of inbreeding and outbreeding. It is therefore of interest to consider the implications of inbreeding for the above results. If selection is weak, the evolutionary dynamics of an inbreeding population can be approximated to the order of second-order terms in *s* by using the neutral value of the within-population inbreeding coefficient, *F*_*IS*_ (Charlesworth, Nordborg & Charlesworth, 1997). The modifications to the relevant equations are given in section 5 of the Supplementary Material, equations (S13) – (S17).

Only the case of a subdivided population will be considered in detail here; the results for large *M* provide a picture of what is expected for a single large population. If *B*_*l*_ for a single locus in a partially inbreeding population is defined as the difference between the mean fitness of random matings within demes and the mean fitness of fully homozygous individuals formed by sampling alleles within demes, and *H*_*l*_ as the difference in mean fitness between random matings between demes and random matings within demes, no changes to equations (6) and (7) are needed.

Table 2 shows results for *N*_*e*_ = 100 and *F*_*IS*_ = 0.9, which can be compared with the results for the same values of *N*_*e*_ and the selection, mutation and migration parameters in Table 1. (*N*_*e*_ with inbreeding is obtained by dividing *N*_*e*_ for the corresponding random mating case by (1 + *F*_*IS*_) (Laporte & Charlesworth, 2002; Pollak, 1987), so that this example corresponds to a larger actual deme size than in Table 1.) It has long been known that the greater effectiveness of selection on a single locus with inbreeding causes the genetic load due to selection against non-recessive deleterious mutations to approach *u* instead of 2*u* in a large population when mutation rates are low (Charlesworth & Charlesworth, 1987; Crow 1970; Lande & Schemske. 1985). The mean genetic load in an inbreeding metapopulation is thus expected to be reduced by inbreeding (Glémin *et al.*, 2003; Roze & Rousset, 2004), as can be seen by comparing Tables 1 and 2. With *M* = 10, both the exact and approximate values of the mean load approach *u*, even with the high mutation rate, and are substantially less than the values for the corresponding random mating cases, including the examples with lower *M* values.

**Table 2.**
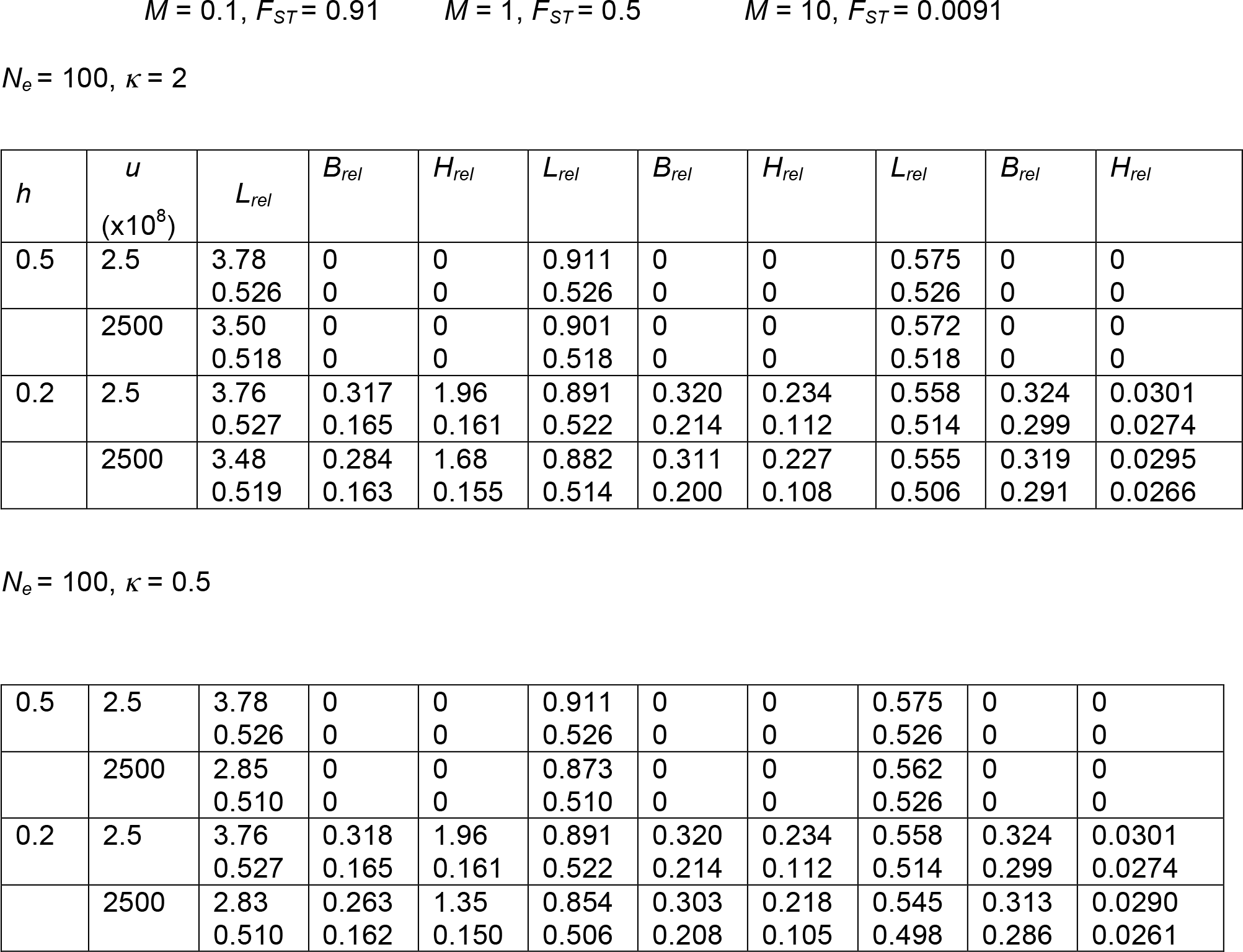
Genetic loads and heterosis in a subdivided population with inbreeding (*F*_*IS*_ = 0.9) All results are expressed relative to the diploid mutation rate, *2u*, as indicated by the subscript *rel*. The expected loads are denoted by *L*_*rel*_.The inbreeding depression values (*B*_*rel*_) were calculated from equation (6), and the heterosis values (*H*_*rel*_) from equation (7). The selection coefficient *s* was 2.5 × 10^−3^. The upper items in the cells are the exact expectations over the probability distribution of local allele frequencies, using equations (S13) and (S14), where *α* and *β* in the latter are replaced by *α* + *Mq*_*t*_ and *β* + *Mp*_*t*_, respectively. The lower items in the cells were calculated from the linearized approximation to this distribution, using equations (S11) and (S12).

| $h$ | $u$<br>($\times 10^8$) | $L_{rel}$ | $B_{rel}$ | $H_{rel}$ | $L_{rel}$ | $B_{rel}$ | $H_{rel}$ | $L_{rel}$ | $B_{rel}$ | $H_{rel}$ |
| --- | --- | --- | --- | --- | --- | --- | --- | --- | --- | --- |
| 0.5 | 2.5 | 3.78 | 0 | 0 | 0.911 | 0 | 0 | 0.575 | 0 | 0 |
|  |  | 0.526 | 0 | 0 | 0.526 | 0 | 0 | 0.526 | 0 | 0 |
|  | 2500 | 3.50 | 0 | 0 | 0.901 | 0 | 0 | 0.572 | 0 | 0 |
|  |  | 0.518 | 0 | 0 | 0.518 | 0 | 0 | 0.518 | 0 | 0 |
| 0.2 | 2.5 | 3.76 | 0.317 | 1.96 | 0.891 | 0.320 | 0.234 | 0.558 | 0.324 | 0.0301 |
|  |  | 0.527 | 0.165 | 0.161 | 0.522 | 0.214 | 0.112 | 0.514 | 0.299 | 0.0274 |
|  | 2500 | 3.48 | 0.284 | 1.68 | 0.882 | 0.311 | 0.227 | 0.555 | 0.319 | 0.0295 |
|  |  | 0.519 | 0.163 | 0.155 | 0.514 | 0.200 | 0.108 | 0.506 | 0.291 | 0.0266 |
| $$N_e = 100, \kappa = 0.5$$ |  |  |  |  |  |  |  |  |  |  |
| 0.5 | 2.5 | 3.78 | 0 | 0 | 0.911 | 0 | 0 | 0.575 | 0 | 0 |
|  |  | 0.526 | 0 | 0 | 0.526 | 0 | 0 | 0.526 | 0 | 0 |
|  | 2500 | 2.85 | 0 | 0 | 0.873 | 0 | 0 | 0.562 | 0 | 0 |
|  |  | 0.510 | 0 | 0 | 0.510 | 0 | 0 | 0.526 | 0 | 0 |
| 0.2 | 2.5 | 3.76 | 0.318 | 1.96 | 0.891 | 0.320 | 0.234 | 0.558 | 0.324 | 0.0301 |
|  |  | 0.527 | 0.165 | 0.161 | 0.522 | 0.214 | 0.112 | 0.514 | 0.299 | 0.0274 |
|  | 2500 | 2.83 | 0.263 | 1.35 | 0.854 | 0.303 | 0.218 | 0.545 | 0.313 | 0.0290 |
|  |  | 0.510 | 0.162 | 0.150 | 0.506 | 0.208 | 0.105 | 0.498 | 0.286 | 0.0261 |

For *B*_*rel*_ and *H*_*rel*_, the patterns are similar to those found with random mating but with substantially smaller values, except that the exact value of *B*_*rel*_ with *h* = 0.2 does not change greatly with changes in *M*; instead, it remains quite close to the value given by substituting the deterministic equilibrium value of *q* from equation (S15) into equation (6) with *F*_*IS*_ = 0 (approximately 0.33, as opposed to 1.5 with random mating). This appears to be due to a sharp increase in the mean of *q* as *M* decreases, which compensates for the decline in 1 − *F*_*s*_. Consistent with this interpretation, the linearized approximations for *B*_*rel*_ and *H*_*rel*_ perform poorly with *M* < 10. The approximate constancy of *B*_*rel*_ with changing *M* is also found with *N*_*e*_ = 1000 and *h* = 0.2. In this case, the mean allele frequency changes little with *M*, while *F*_*s*_ is held to a low level by selection, and the linearized approximation works well for all *M* values; *H*_*rel*_ is always small (approximately 0.05 for *M* = 0.1, and 0.02 for *M* = 1).

## 3 EMPIRICAL EVIDENCE ON INBREEDING DEPRESSION AND HETEROSIS IN SUBDIVIDED POPULATIONS

### 3.1 Plant populations

Table S2 of the Supplementary Material shows some empirical estimates of the mean inbreeding depression and heterosis for flowering plants with subdivided populations. The studies concerned were selected on the basis that estimates of lifetime fitness were available, rather than single fitness components, and that data on multiple local populations were collected. Overall, both *B* and *H* are usually modest in size. The largest *B* values in the table are 2.52 ± 0.60 and 3.32 ± 0.82 (where ± indicates the standard error), for self-incompatible populations of *Arabidopsis lyrata* (Oakley, Spoelhof & Schemske, 2015) and predominantly outcrossing populations of *Sabatia angularis* (Spigler, Theodorou & Chang, 2017), respectively. These values are considerably larger than the value of 1.5 expected with *h* = 0.2 and *U* = 1 for deterministic equilibrium and outcrossing, but the large standard errors means that it is not possible to tell whether they are too large to be explained on this basis. Strongly selected deleterious mutations with small *h* values contribute substantially to inbreeding depression in large outcrossing populations (Charlesworth & Charlesworth, 1987; Simmons & Crow, 1977), although they are likely to be purged from highly inbred or very small populations (Lande & Schemske, 1985; Wang *et al.*, 1999). They will not, however, contribute much to heterosis between populations, since their frequencies are not greatly affected by drift (see section 4.3).

The largest values of *H* in the table are 1.20 ± 0.47 for small populations of *Hypericum cumulicola* (Oakley & Winn, 2012) and 1.43 ± 0.47 for partially selfing populations of *Arabidopsis lyrata* (Oakley, *et al.*, 2015). *F*_*ST*_ among the small populations of *H. cumulicola* was 0.74 (from Table S1 of Oakley & Winn [2012]), and 0.78 among partially selfing populations of *A. lyrata* (Mable & Adam, 2007). Patterns of variability at molecular markers indicate that both sets of populations are partially inbred (mean *F*_*IS*_ of 0.74 and 0.44 for small populations of *H. cumulicola* and partially selfing *A. lyrata*, respectively).

### 3.2 Animal populations

Relevant data where both *B* and *H* were measured for net fitness are less abundant for animal populations, despite the fact that studies showing heterosis in interpopulation crosses were pioneered in *Drosophila pseudoobscura*, using data on recessive lethals (Wright, Dobzhansky & Hovanitz, 1942) and several fitness components (Vetukhiv, 1953, 1956, 1957). However, outbreeding depression, reflecting the accumulation of Dobzhansky-Muller genetic incompatibilities between local populations, is often observed in animals, especially in hermaphrodite species with high frequencies of self-fertilization (Dolgin, Charlesworth, Baird & Cutter, 2007; Escobar *et al.*, 2008). This makes it hard to assess the extent of heterosis.

The most informative study is probably that of Lohr & Haag (2015), on groups of populations of *Daphnia magna* inhabiting both small and large ponds. Their results on the net reproductive output per individual under laboratory conditions provide a measure of net fitness. The small ponds had *B* and *H* values of 0.13 ± 0.38 and 0.96 ± 0.26, respectively; the corresponding values for large ponds were 1.69 ± 0.31 and – 0.30 ± 0.30. Previous work showed that *F*_*ST*_ for molecular markers was considerably higher among small than large populations (0.70 versus 0.38 for microsatellites), with much lower within-population diversity in the small populations (Walser & Haag, 2012).

### 3.3 Small versus large populations

Several of the plant studies also allowed groups of local populations with small census sizes to be contrasted with groups with larger size, or partially self-fertilizing populations to be contrasted with outcrossing populations. If smaller population size and a greater extent of selfing are both associated with reduced *N*_*e*_ and *M*, as is likely to be the case (Charlesworth, 2003), *H* should be larger for the small/partially inbreeding populations than the large/outcrossing ones, with the opposite pattern for *B.* A difficulty with this prediction is that increased inbreeding, with its purging effects, causes a reduction in *H* as well as *B*, which could dilute or even reverse the effect of inbreeding on *N*_*e*_ and *M*. However, even the self-incompatible populations of *A. lyrata* have substantial *F*_*IS*_ values, with a mean of 0.20 (Mable & Adam, 2007), so this difficulty is probably not as severe as might have been expected.

If the *Daphnia* results are included, and only the Willi (2013) study is used for *A. lyrata* (to avoid pseudoreplication), five independent contrasts are available for *H* and four for *B*. All five of the *H* differences are in the direction of a greater value for small/selfing populations, with a mean difference of 0.70 ± 0.23 (*P* = 0.031 on a sign test, or 0.019 on a one-tailed paired *t*-test, *t* = 3.06). But of the four contrasts for *B*, one has a greater value for small/selfing populations (*A. lyrata*), and the mean contrast between large/outcrossing and small/selfing populations is 0.70 ± 0.42, *t* = 1.65, *P* > 0.10. Given that several of these studies reported significant differences in their measures of heterosis between small/selfing and large/outcrossing populations, it seems fairly clear that there is indeed a significant effect for *H* in this direction; the situation with *B* is less clear. This is perhaps not unexpected, since the relative effects of differences in *M* and *N*_*e*_ are more marked for *H* than *B* (Tables 1 and 2).

A similar question can be asked about the genetic load. While it is not feasible to estimate the absolute value of the load, because the fitness of the hypothetical mutation-free genotype is unknown, it is possible to compare the mean fitnesses of small and large populations, or selfing and outcrossing populations. The natural logarithm of the ratio of mean fitnesses reflects the difference in the sums of the *L*_*l*_ over all loci between the two groups, under the multiplicative fitness assumption. In the studies listed in Table S2, and in many other studies (Leimu *et al.*, 2006), the individuals whose fitnesses are estimated were the product of random matings among parents sampled from natural populations, and so did not necessarily represent the fitnesses of individuals produced by the natural mating system. This means that the corresponding load for a single locus is described by equation (5), with the proviso that the mean *q* is the value expected under the natural mating system, not random mating.

The expectation that the load is lower for large than small populations holds true for this measure, consistent with the results of a large meta-analysis (Leimu *et al.*, 2006). The expectation for a comparison between selfing and outcrossing populations is less clear, since selfing is expected to reduce the mutational load caused by partially recessive mutations (Charlesworth & Charlesworth, 1987; Lande & Schemske, 1985), whereas increased drift due to smaller effective population size and reduced migration has the opposite effect. For the cases in Table S2, a significantly positive correlation between population size and fitness (*P* < 0.05) was found by Paland & Schmid (2003); for the data of Oakley & Winn (2012) on *H. cumulicola* and Lohr & Haag (2015) on *D. magna*, the differences in log mean fitness between large and small populations were 1.13 ± 0.51 and 1.22 ± 0.31, respectively, indicating a clear effect of population size in the expected direction.

For the outcrossing versus selfing comparisons in Table S2, the corresponding load differences were 0.50 ± 0.50 (Busch, 2006), 1.25 ± 0.29 (Willi, 2013), and 0.96 ± 0.53 (Oakley *et al.*, 2015) (the last two values are both for *A. lyrata*). This suggests that the effect of differences in levels of inbreeding between the outcrossing and selfing populations, which are relatively modest in the case of *A. lyrata*, are outweighed by difference in effective deme size and/or level of population subdivision.

## 4 DISCUSSION

### 4.1 The relation between the mutation rate and the genetic load

The theoretical results described in section 2 show that, when the scaled selection coefficient (*γ* = 2*N*_*e*_*s*) is ≫ 1 in a single population or in an island metapopulation
with a scaled migration rate *M* = 4*N*_*e*_*m* ≫ 1, the expected genetic load for a single locus at mutation-selection-drift equilibrium is close to the deterministic value, consistent with previous studies that assumed low mutation rates, e.g. Kimura *et al.* (1963). The load strongly depends on the mutation rate even when the classic Haldane (1937) formula (*L*_*l*_ = 2*u*) is not exact.

Dr Frank Johannes has suggested to me that a high rate of reversion from deleterious to beneficial alleles may result in only a weak dependence of the load on the mutation rate, but this does not appear to be case when the results with *κ* = 2 are compared with those with *κ* = 0.5 in Tables 1, 2 and S1. Although a high rate of reversion from mutant to wild-type somewhat reduces the load, it does not abolish its dependence on the mutation rate (see the parts of the Table with *γ* ≫ 1). In this area of parameter space, selection to reduce the mutation rate would be expected if *M* or *N*_*e*_, as well as the total genomic mutation rate, are sufficiently large (Drake, Charlesworth, Charlesworth & Crow 1998; Sniegowski, 2000; Lynch, Ackerman, Gout, Long, Sung, Thomas & Foster, 2016). This follows from the fact that, in a randomly mating population with free recombination, the selection coefficient on a modifier of the mutation rate is approximately equal to the product of the mean selection coefficient on the deleterious mutations themselves, and the change in the genome-wide deleterious mutation rate (δ*U*) caused by the modifier. In a completely selfing population, the selection coefficient is approximately equal to 0.5 δ*U* (Drake *et al.,* 1998). δ*U* will tend to be larger, the larger *U,* leading to stronger selection on the mutation rate.

The picture is very different for the stochastic regime with *γ* = 2*N*_*e*_*s* ≤ 1, or *M* < 1 where the fact that the deleterious allele has an appreciable probability of being at a high frequency means that the load is largely determined by the strength of selection, and is only weakly affected by *U* (see the parts of the tables with *γ* ≤ 1). Here, selection on modifiers that reduce the mutation rate is likely to be ineffective in the face of drift (Lynch *et al.*, 2016).

These considerations suggest that the methylation status of CG sites in *A. thaliana,* with its high epimutation rate (Van De Graaf *et al.*, 2015), is probably close to being selectively neutral with respect to purifying selection at most such sites, otherwise selection should have reduced the mutation rate to a much lower level. This is consistent with the results of a population survey where the site frequency spectrum of CG epialleles was used to estimate the intensity of selection, and no significant departure from neutrality was detected (Vidalis *et al.*, 2016). It is, of course, possible that a subset of these sites are functionally significant, and contribute to genetic load and interpopulation heterosis.

Another possibility is that the functional consequences of these epimutations relate to quantitative traits under stabilizing selection; there is experimental evidence that quantitative trait variation in *A. thaliana* can be caused by epimutations (Quadrana & Colot, 2016). In this case, the relevant contribution to the genetic load would come from the epigenetic component of the genetic variance in the trait, multiplied by the factor *S* that determines the strength of selection on the trait as a whole – see equations (2) of Charlesworth (2013). Under a balance between stabilizing selection and mutation, the magnitude of this component in a highly selfing species like *A. thaliana* depends on the mutational variance, *V*_*m*_ (Charlesworth & Charlesworth, 1995; Lande & Porcher, 2015). Empirical estimates of *V*_*m*_ for a range of quantitative traits are available from the literature (Halligan & Keightley, 2009); these include potential contributions from epigenetic as well as genetic variants. These estimates are surprisingly high compared with what might be expected from DNA sequence mutation rates if each trait were controlled by independent sets of genes (Johnson & Barton, 2005), implying either a high degree of pleiotropy across traits or a substantial contribution from epialleles to quantitative trait variability. Further empirical research is needed to discriminate among these hypotheses.

### 4.2 General patterns expected from the theoretical predictions

Equations (6) and (7), which apply to both randomly mating and partially inbreeding populations, bring out the complementary relations between the measures of inbreeding depression and heterosis in a metapopulation. For a given mean frequency of deleterious alleles in the metapopulation, increased differentiation among demes as measured by *F*_*s*_ increases *H*_*l*_ and reduces *B*_*l*_, provided that the dominance coefficient *h* is less than one-half. An increase in mean *q* due to drift tends to increase both *H*_*l*_ and *B*_*l*_, as long as it remains less than one-half. If deme sizes or migration rates are so small that mean *q* is much greater than one-half, *H*_*l*_ and *B*_*l*_ can be reduced compared with large *N*_*e*_*m* values; this is more likely to affect *B*_*l*_, since the associated increase in *F*_*s*_ works against *B*_*l*_ but in favor of *H*_*l*_. With semidominance (*h* = ½), the mean load given by equation (5) (and its modification for an inbreeding population, equation S17) is independent of *F*_*s*_, and always increases with mean *q*. With *h* < ½, the mean load increases with both mean *q* and *F*_*s*_.

The numerical results described in sections 2.4 and 2.5, where the load parameters are expressed relative to the diploid mutation rate (Tables 1 and 2), suggest that the main effect of *N*_*e*_*m* on these measures for weakly selected mutations is due to changes in *F*_*s*_ rather than mean *q*, except when *M* ≤ 1, since the predictions from the linearized approximation for *F*_*s*_ (equation S12), which assumes that mean *q* is equal to the deterministic equilibrium value *q*\*, perform quite well for *M* > 1 unless the mutation rate is very high.

However, unless *h* approaches zero, *L*_*rel*_ > 1 and *B*_*rel*_ ≫ 1 require an increase in mean *q* above *q*\*, unless *h* and *F*_*IS*_ are close to zero. Even with small *h*, it is likely that *H*_*rel*_ remains < 1 without an increase in mean *q*, as described in section 5 of the Supplementary Material. As expected from these considerations, and in line with previous theoretical results (Glémin *et al.*, 2003; Roze & Rousset, 2004), Table 1 shows that smaller *N*_*e*_ and *M* generally lead to larger *L*_*rel*_ and *H*_*rel*_, and smaller *B*_*rel*_. Inbreeding within demes leads to lower values of *L*_*rel*_, *H*_*rel*_ and *B*_*rel*_ compared to the same *N*_*e*_ and *M* with random mating.

### 4.3 Relations between theoretical predictions and empirical results

On the assumption of multiplicative fitnesses, the values of the load, inbreeding depression and heterosis contributed by deleterious mutations across the whole genome can be roughly predicted by the products *U L*_*rel*_, *U B*_*rel*_ and *U H*_*rel*_ where *U* is the total diploid genomic mutation rate to deleterious mutations. For *Drosophila*, data on DNA sequence mutation rates and levels of selective constraint imply that *U* for weakly selected mutations is approximately one (Keightley, 2012; Charlesworth, 2015). Given that the number of genes in the genomes of flowering plants such as *Arabidopsis* is approximately twice as large as in *Drosophila* (*www.biology-pages.info/G/GenomSizes.html*) and that the mutation rates per basepair in *A. thaliana* and *D. melanogaster* are similar (Keightley *et al.*, 2014; Ossowski *et al.*, 2010), it seems safe to assume that *U* for plants like *Arabidopsis* is at least one, and may be much larger. The results in Tables 1 and 2 should thus provide a rough guide as to what parameter values to expect for natural populations of flowering plants, when contrasting populations with complete outbreeding with those with a high level of inbreeding. The empirical results summarised in Table S2 are broadly consistent with the range of parameter values in these tables for *U* of order 1.

The procedures described in sections 2 and 3 of the Supplementary Material can be use to assess the extent to which the data on small and large demes can be explained by the mutation-selection-drift model. Predictions were generated for a wide range of selection coefficients, from *s* = 2.5 × 10^−5^ to *s* = 0.05 and for *h* = 0.2, which cover the range for deleterious nonsynonymous mutations estimated from population genomics data. These were applied to the infinite island model, using estimates of *F*_*IS*_ and *F*_*ST*_ from molecular marker data for large and small populations of *Hypericum cumulicola* (Oakley & Winn, 2012), and *F*_*ST*_ for *Daphnia magna,* where there is no statistically significant evidence for local inbreeding (Lohr & Haag, 2015). The scaled migration rate, *M*, was estimated from (1 − *F*_*ST*_)/*F*_*ST*_. The *N*_*e*_ values for small and large populations were arbitrarily set to 100 and 1000, respectively.

Figure 1 shows the resulting plots of the predicted difference in mean relative load between small and large populations (Δ*L*_*rel*_) against *s,* as well as the predicted *B*_*rel*_ and *H*_*rel*_ values for small and large populations. Overall, the predictions are not very sensitive to *s* until it becomes very large compared with the mean value suggested by population genomics analyses of *Drosophila*, which is approximately 10^−3^ for nonsynonymous mutations (Campos *et al.*, 2017), in which case both Δ*L*_*rel*_ and *H*_*rel*_ start to decline rapidly.

**Figure 1.**
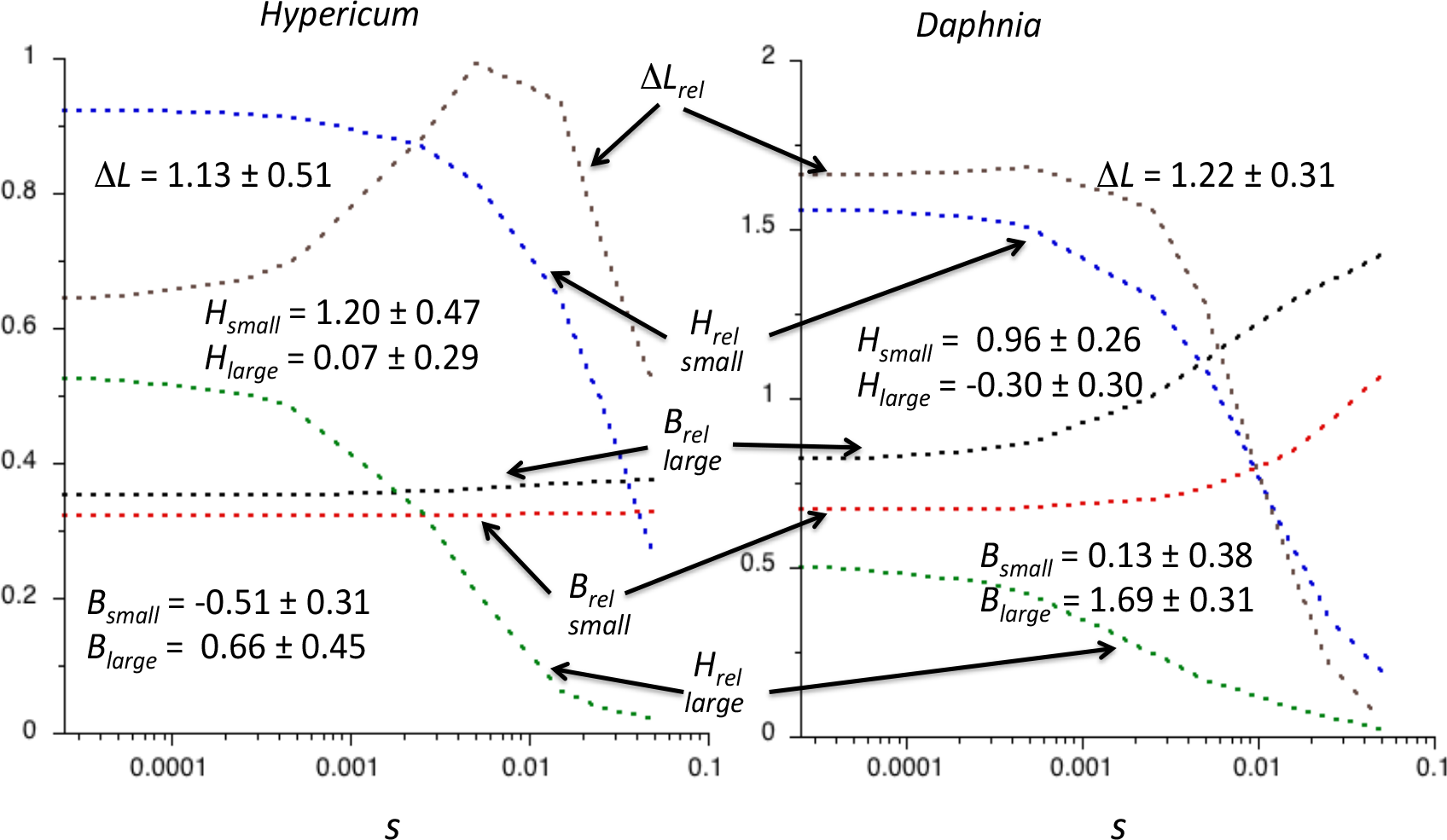
Plots against *s* of the predicted values of *B*_*rel*_ (red and black curves, for small and large populations, respectively) *H*_*rel*_ (blue and green curves, for small and large populations, respectively) and Δ*L*_*rel*_ (grey curves) for the *H. cumulicola* (left) and *D. magna* (right) parameters described in the text. The empirical estimates of these parameters and their standard errors are shown as inserts inside the panels.

In these two examples, Δ*L*_*rel*_ with *s* = 5 × 10^−4^ was 1.07 for *H. cumulicola* and 1.68 for *D. magna*, with empirical values of Δ*L* of 1.13 ± 0.51 for *H. cumulicola* and 1.22 ± 0.31 for *D. magna*. Similar results apply to *H*_*rel*_ (the differences between small and large populations were 0.41 for *H. cumulicola* and 1.08 for *D. magna*), compared with observed *H* values of 0.50 ± 0.55 and 1.26 ± 0.39, respectively. However, the magnitudes of the observed differences in *B* between small populations and large populations of *H. cumulicola* (− 1.17 ± 0.55) and for *D. magna* (− 1.55 ± 0.49) are much larger than those of the differences in *B*_*rel*_ ( − 0.03 for *H. cumulicola* and – 0.19 for *D. magna*). This reflects a large (but non-significantly) negative *B* value for the small populations of *H. cumulicola* and a near zero value for small populations of *D. magna*, compared with *B*_*rel*_ values of 0.32 and 0.68 for the respective small populations. The theoretical results may thus overestimate *B* for small populations, possibly because the island model does not capture bottleneck effects associated with extinction-recolonization events (see section 4.5).

Figure 2 shows the dependence of the results on the dominance coefficient for examples with weak and strong selection, using the demographic parameters for *Daphnia*, since dominance will have the greatest effect when there is no inbreeding. It can been see that a high degree of recessivity is necessary to obtain large *H*_*rel*_ values when selection is strong. With weak selection, there is a nearly linear dependence of *B*_*rel*_ and *H*_*rel*_ on *h*, whereas Δ*L*_*rel*_ is almost independent of *h*. With strong selection, Δ*L*_*rel*_ is always close to zero, and in fact is slightly negative for *h* values close to 0.1, reflecting the effect of purging.

**Figure 2.**
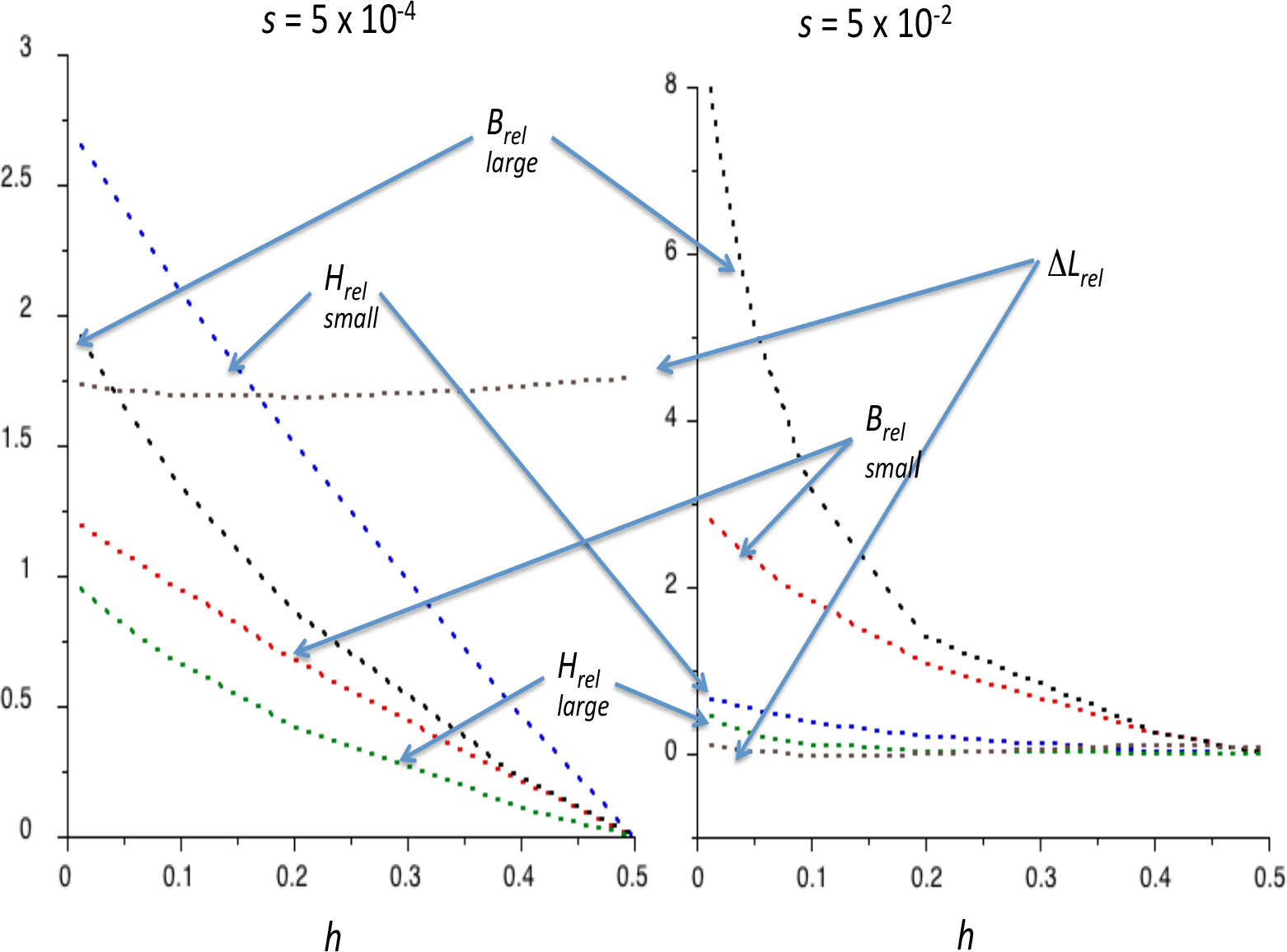
Plots against *h* of the predicted values of *B*_*rel*_ (red and black curves, for small and large populations, respectively) *H*_*rel*_ (blue and green curves, for small and large populations, respectively) and Δ*L*_*rel*_ (grey curves) for weak selection (left) and strong selection (right), using the *D. magna* demographic parameters described in the text.

The high standard errors of the estimates of *B* and *H*, especially *B*, in many of the studies summarised in Table S1 makes quantitative tests for most individual cases unreliable, especially as estimates of *F*_*ST*_ and *F*_*IS*_ are not always available. The analysis of the coefficients of variation of *L, B* and *H* in section 6 of the Supplementary Material suggests that variance in allele frequencies between demes is not a major source of the large standard errors, which must therefore arise from experimental error and environmental variation. This suggests that an intensive study of a favourable system, resulting in more precise parameter estimates, would provide a rigorous test of the theory. Overall, however, it seems clear that weak selection, of the magnitude of that inferred from population genomics data, together with a substantial degree of recessivity of deleterious mutations, is required to explain the data on Δ*L* and *H*, on the hypothesis of purely mutational load.

### 4.5 Caveats and complications

There are several important caveats concerning attempts to interpret the data in terms of the simple models used here, which mean that the quantitative predictions should be regarded with caution. First, the assumption of statistical equilibrium for drift, mutation and selection under an island model of migration is questionable, since species with small population sizes are likely to be subject to bottlenecks and extinction-recolonization processes. Relatively few attempts to include such complications in models of inbreeding depression and heterosis have been made, with the recent exception of Spigler *et al.* (2017).

It should be noted, however, that a set of populations subject to recent bottlenecks is likely to experience effects that are similar to, but less severe than, making an inbred line, with effects predominantly on *F*_*s*_ rather than mean *q* (Balick, Do, Cassa, Reich & Sunyaev, 2015; Simons, Turchin, Pritchard & Sella, 2014). *B* will thus be greatly reduced, because of loss of variability. *H* will be increased, but only to an extent that corresponds to the inbreeding depression in the ancestral population, since there is no time for an accumulation of deleterious mutations at a high frequency that is responsible for the high values seen for the small *M* values in Tables 1 and 2. There is, therefore, no reason to expect recent bottlenecks to result in much higher *H* values than those described here. Recurrent bottlenecks associated with a long history of extinction and recolonization, with the associated very high *F*_*ST*_ values (Pannell & Charlesworth, 1999; Charlesworth & Charlesworth, 2010, Chap. 7), might have such an effect of *H*, but there is no evidence for extremely high *F*_*ST*_ values in the examples analysed here.

Second, the assumption of independence among loci used here is likely to be violated in finite inbreeding and subdivided populations, due to associations among loci created by linkage disequilibrium and identity disequilibrium, resulting in Hill-Robertson interference among selected loci (Felsenstein, 1974). Only modest effects of such interference among slightly deleterious mutations on levels of inbreeding depression in partially selfing populations have been found for multiplicative fitness models with dominance coefficients of the magnitude assumed here (Bersabé, Caballero, Pérez-Figuereoa, & García-Dorado, 2016; Charlesworth, Morgan & Charlesworth, 1992; Kamran-Disfani & Agrawal, 2014; Roze, 2015), unless the population size is very small. Larger effects of interference may occur for the small dominance coefficients (*h* ≤ 0.02) for severely deleterious or recessive lethal mutation (Kelly, 2007; Lande, 1994; Lande & Porcher, 2017; Porcher & Lande, 2016), leading to much higher frequencies of deleterious mutations than expected in the absence of interference, but the low values of *B* in most studies in Table S2 suggest that the contributions from such mutations are minor. Indeed, they are likely to be purged from partially inbreeding populations, unless mutation rates are very high (Lande & Schemske, 1985; Kelly, 2007). Lack of independence caused by synergistic epistasis among slightly deleterious mutations appears to somewhat reduce the effect of inbreeding on the level of inbreeding depression (Charlesworth, Charlesworth & Morgan, 1991), especially with very high rates of selfing, but its effect on heterosis has not been investigated.

The consequences of interference for *B* and *H* in subdivided populations have only just started to be investigated by rigorous models (Roze, 2015); in this study, interference appeared to have modest effects of increasing *B* and reducing *H*, mainly by reducing *F*_*s*_. However, simulations that incorporate distributions of selection coefficients that are consistent with the population genomics estimates are currently lacking, so that it is currently unclear whether interference alone could explain the discrepancies between observations and theory for *B* that were described above.

Finally, the possible contributions of strongly selected deleterious mutations and variability maintained by heterozygote advantage have been ignored. As shown above, deleterious mutations with selection coefficients much greater than 10^−3^ are unlikely to contribute significantly to *H* and Δ*L* in a subdivided population, unless deme size is extremely small with a very high level of population subdivision. Heterozygote advantage can be studied by a straightforward extensions of the methods employed here, and its consequences for inbreeding depression in a subdivided population has been investigated by Whitlock (2002).

His results are extended to predictions of *H* with random mating within demes in the Supplementary Material, section 8. Equation (S21) shows that loci under balancing selection can contribute to heterosis. With weak selection, *H* is approximately equal to the production of *F*_*ST*_ and the equilibrium genetic load under balancing selection. Current evidence from population genomic surveys suggest that loci maintained by heterozygote advantage and other forms of balancing selection are sparsely distributed across the genome (Charlesworth 2006; Gao, Przeworski & Sella 2015; Siewert & Voight, 2017); the intensity and mode of selection at such loci is largely unknown. Indirect evidence suggests, however, that balancing selection contributes significantly to genetic variation in fitness in *Drosophila* (Charlesworth, 2015) and to inbreeding depression (Charlesworth & Charlesworth, 1999), and so could be partly responsible for the observed values of *B* and *H*.

## Acknowledgements

I thank Diala Abu Aswad and Deborah Charlesworth for their comments on a previous version of this manuscript.

## Data accessibility

No new data were generated for this paper. Computer code and output files will be deposited in Dryad on acceptance.

## Author contributions

Brian Charlesworth designed and conducted the research, and wrote the manuscript.

## Supplementary Material

### 1. Exact and approximate deterministic equations for a randomly mating population

The equilibrium frequency of A_2_, *q*\* satisfies the following cubic equation (Bürger, 2000, p.100):

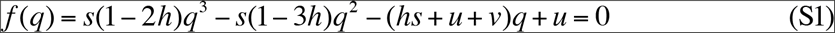

While a formal solution to equation (S1) can be written down, in practice it is simpler to solve it numerically by Newton-Raphson iteration, although it reduces to a quadratic equation with an explicit solution when *h* = 0.5. The exact value of the equilibrium genetic load, *L*\*, can be obtained by substituting the value of *q*\* obtained in this way into equation (S1).

If *t* = *hs* ≫ *u*, second and third-order terms in *q*\* can be ignored and we obtain:

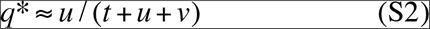

The corresponding approximate equilibrium genetic load is obtained by ignoring second-order terms in *q*\* in equation (S1), yielding:

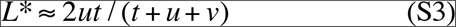

*L*\* is always smaller than the classical approximate value of 2*u* for non-recessive autosomal mutations (Haldane 1937), but approaches 2*u* when *t* ≫ (*u* + *v*). If we write *u* = *κv*, where *κ* measures the mutational bias towards or against deleterious alleles it is easily seen from equation (S3) that the derivative of *L*\* with respect to *v* is proportional to *κt*, so that the equilibrium load increases with the mutation rate, although the dependence is weak if *κ* ≪ 1, i.e. when mutation is strongly biased towards the beneficial allele.

If *u* + *v* approaches *t*, this approximation breaks down, and the full solution to equation (S1) needs to be used. Table S1 and Figure S1 illustrate the dependence of the equilibrium load on the selection and mutation parameters for such cases. The main features of the results are as follows. The ratio of the *L*\* to the classical value of 2*u*, *L*_*rel*_, decreases with the mutation rate. A high rate of reversion from the deleterious mutant state to wild-type, and a small dominance coefficient (*h*), enhances this effect. Nonetheless, even when *u* is comparable to *t*, *L*\* is always several times greater than with *u* ≪ *t*, although its relation to *u* is one of diminishing returns. For example, for *h* = 0.2 and *κ* = 0.5 (the situation in Figure S1 where the mutation rate has the least effect on the load), *L*_*rel*_ = 0.983 for *u*/*s* = 0.001; with *u*/*s* = 1 (1000 times larger), *L*_*rel*_ = 0.0894, representing an absolute load that is 91 times greater. Even with a much more extreme mutational bias towards the beneficial alleles of *κ* = 0.1, the ratio of *L* values for these two values of *u*/*s* is 23.

### 2. Exact stochastic equations for a randomly mating population

For computational purposes, it is preferable to write the relative fitnesses of A_1_A_1_, A_1_A_2_ and A_2_A_2_ as 1 + *s´*, 1 + *h´s´* and 1, respectively (Charlesworth & Jain, 2014), where *s´* = 1/(1 − *s*) and *h´* = 1 − *h*. The probability density of *p* = 1 − *q* is then an increasing exponential function of *p* and *s´*, provided that *h* > ½. For small *s*, such that *O*(*s*^2^) terms can be ignored, as assumed here, *s´* can be equated to *s* to a good level of approximation. From the general expression for the stationary distribution of allele frequencies (Wright, 1937; Charlesworth & Charlesworth, 2010, Chap. 5), the probability density of *p* is given by:

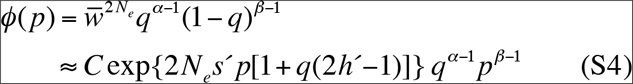

Writing *γ* = 4*N*_*e*_*s´*, the exponential term in equation (S4) can be expanded as:

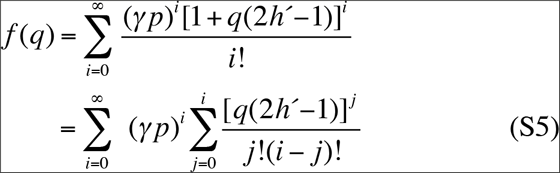

Substituting the double series into equation (S4) and integrating between 0 and 1, we obtain:

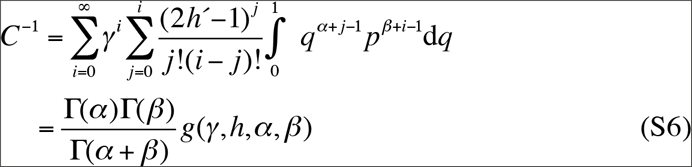

where:

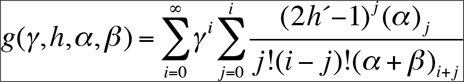

and (*x*)_*i+j*_ is the Pochhammer symbol, such that (*x*)_0_ = 1, (*x*)_1_ = *x*, (*x*)_*k*_ = *x*(*x* + 1)… (*x* + k −1).

This approach provides a compact expression for the moments (*μ*_*k*_) about zero, such the expectation of *q*^*k*^ is given by:

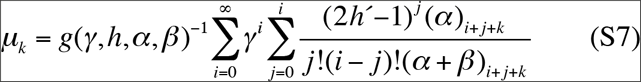

The expected genetic load is given by:

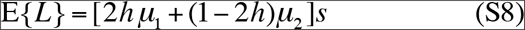

In the case of semidominance (*h* = ½), equations (S6) and (S7) simplify, since the only term in the sum over *j* is for *j* = 0. This yields expressions involving the confluent hypergeometric function (Kimura, Maruyama & Crow, 1963; Charlesworth & Jain, 2014). In this case, for *β* ≪ 1 and *γ* ≫ 1, the expected value of *q* is close to the deterministic value 2*u*/*s* and the expected load approaches 2*u*. For larger values of *β*, this is no longer true and the dependence of the load on the mutation and selection parameters must be evaluated numerically.

A computer program for carrying out these evaluations has been developed. The powers of *γ* and 2*h*´ − 1 in equations (S5) and (S6) are both positive when *s*´ > 0 and *h* < ½, so that logarithms of the terms in the series and their partial sums can be obtained, avoiding overflow problems when *γ* is ≫ 1. If the *i*th term in the series is *X*_*i*_, and the corresponding partial sum is *S*_*i*_, we have:

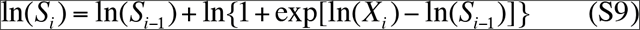

This relation enables the logarithms of the constant *C* and the moments of the distribution to be obtained without difficulty, even when *γ* is as large as 100, although convergence of the series can be slow.

### 3. Approximate stochastic results for a randomly mating population

An approximate analytic expression for the expected load when *t* > *u*, *v* can be obtained by linearizing Δ*q* around the deterministic equilibrium value *q*\* given by equation (S1), in which case *ϕ(q*) is a beta distribution, e.g. Malécot (1969, p.58), Bataillon & Kirkpatrick (2000) and Charlesworth & Charlesworth (2010, p.355).

The inbreeding coefficient *F* corresponding to the variance in *q* is then given by:

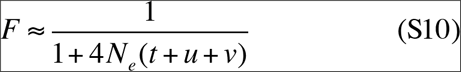

and

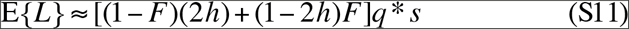

This approximation is valid when *γ* is sufficiently large that third and higher order values of *q* − *q*\* can be neglected, or when *γ* is so small that *φ(q*) approaches the beta distribution for the neutral case.

### 4. Infinite population island model

To deal with the infinite population island model described by equation (8), the above equations are modified by replacing *α* with *α* + *Mq*_*t*_ and *β* with *β* + *Mp*_*t*_, where *q*_*t*_ is the value of *q* for the whole metapopulation and *p*_*t*_ = 1 − *q*_*t*_. To obtain the value of *q*_*t*_ given values of the other parameters, we can iterate the values of *q*_*t*_ in the equation for the first moment of the distribution, given by the equivalent of equation (S4), until *q*_*t*_ and the mean value of *q* converge. This was done using a program for a grid search using the fitness representation 1+ *s*´, 1 + *h*´*s*´, 1, which was iterated for at least 10 cycles with increasingly finer subdivision of the interval (0, 1), or until the proportional difference between successive values of *q*_*t*_ became less than 10^−5^. By using this value of *q*_*t*_ in the series for the second moment of the distribution, equation (S7) can be used to calculate the expected genetic load.

This can be compared to the approximate value obtained by linearization of Δ*q* about the deterministic equilibrium frequency given by equation (S1), using equations (S10) and (S11). The only difference is that *F* is replaced by:

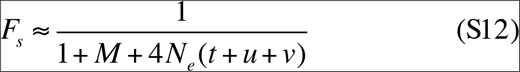

### 5. Subdivided populations with inbreeding

The mean fitness of the population with frequency *p* of allele A_1_, under weak selection with inbreeding coefficient *F*_*IS*_ and relative fitnesses of A_1_A_1_, A_1_A_2_ and A_2_A_2_ of 1 + *s´*, 1 + *h´s´* and 1, respectively, can be written as:

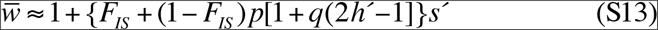

To obtain the equivalent of equation (S4), the term in *F*_*IS*_ in this expression should be given a weighting of 2, corresponding to its contribution to the change in allele frequency due to selection that is obtained by differentiation of the fitness potential function with respect to *p* (Wright, 1969, p.244). The resulting expression for the probability density of *p* is:

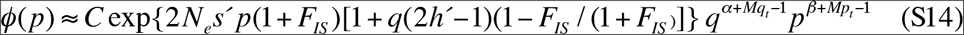

The modifications to equations (S5) – (S7) as far as the selection term is concerned simply require *γ* to be replaced by 2*N*_*e*_(1 + *F*_*IS*_)^−1^*s* and (2*h*´− 1) with (2*h*´− 1)(1 – *F*_*IS*_)/(1 + *F*_*IS*_). The computations of the required moments can thus be carried out by a simple modification of the program used for random mating.

Using the alternative fitness representation of 1, 1 − *hs* and 1 − *s*, the exact equation for the deterministic equilibrium allele frequency *q*\* is:

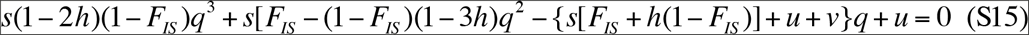

and the equivalent of the approximate equation (S3) is:

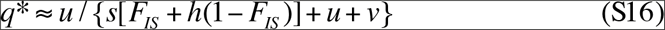

As before, the expected deterministic and stochastic loads can be obtained by substituting the values of *q*\* and the first and second moments of *q* into the relevant expressions for mean fitness. For a population with allele frequency *q*, the load is given by:

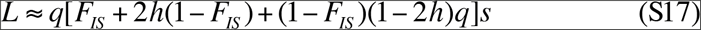

### 6. Upper limits for *L*_*rel*_, *B*_*rel*_ and *H*_*rel*_

The value of *L*_*l*_ at deterministic equilibrium is approximately (1 − *F*_*IS*_/2), if *q* * ≪ 1. Similarly, *B*_*rel*_ with *q* * ≪ 1 is approximately (½ − *h*)/[(*F*_*IS*_ + (1 − *F*_*IS*_)*h*], and *H*_*rel*_ = *F*_*s*_*B*_*rel*_/(1 − *F*_*s*_). For an outcrossing population with the parameters of Table 1, *L*_*rel*_ = 1; with *F*_*IS*_ = 0.9 (Table 2), *L*_*rel*_ = 0.550. The corresponding values of *B*_*rel*_ are 1.5 and 0.326. According to equations (6) and (7), with *h* < ½, 0 < *F*_*s*_ < 1 and mean *q* ≪ 1, both *B*_*rel*_ and *H*_*rel*_ will be less than the deterministic value of *B*_*rel*_. If mean *q* is held constant at *q*\*, equation (5) implies that *L*_*rel*_ increases linearly with *F*_*s*_ towards a maximum of 2.5 with *h* = 0.2 for an outcrossing population, but remains unchanged at 0.5 for a fully inbreeding population: this is because *L*_*rel*_ for a given value of *F*_*s*_ is equal to the product of (1 − *F*_*IS*_)*F*_*s*_ and the deterministic equilibrium value of *B*_*rel*_, plus *L*_*rel*_ at *F*_*s*_ = 0.

### 7. Variances of load, inbreeding depression and heterosis due to inter-deme variation in allele frequencies

From equations (5)-(7), it can be seen that the variances of the load, inbreeding depression and heterosis due to variance in allele frequencies between populations are determined by the 1^st^ through 4^th^ moments of *q*, *μ*_1_ to *μ*_4_. Elementary algebra using these expressions yields the following expressions for these variances for a single locus:

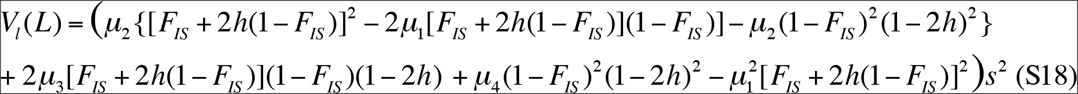

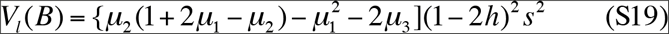

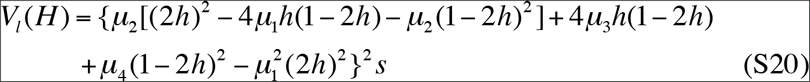

By combining these expressions with the expectations given by equations (5)-(7), the corresponding coefficients of variation (CVs) can be obtained. Numerical values of the CVs can be found using the computational procedures described in sections 2 and 4 above; these also apply to the values of *L* etc when divided by 2*u*. The CVs are generally very large, reflecting the skew of the distribution of allele frequencies towards low frequencies of the deleterious alleles. For example, with the population size, selection and mutation parameters in the upper part of Table 1, the CVs of *L* with *M* = 0.1, 1 and 10 are 72.2, 115 and 49.7; the corresponding values for *B* are 88.4, 79.2 and 41; and those for *H* are 127, 282 and 422.

However, if there are many independent sites that contribute to *L*, *B* and *H*, as is almost certainly the case, their CVs are of the order of the typical individual locus value, divided by the square root of the number of sites, *n*. This can easily be seen in the case when each site has an equal effect, in which case the standard deviation is equal to the product of √*n* and the single locus standard deviation, whereas the expectation is equal to the product of *n* and the single locus expectation. There are likely to be of the order of 20,000 genes or more in a plant genome, with a mean of 1500 coding nucleotide sites per gene, 70% of which can generate nonsynonymous mutations, giving *n* = 2.1 × 10^7^, and √*n* = 4582. Even with a single locus CV of 500, the CV for the trait as a whole is only 0.11 in this case.

### 8. Heterozygote advantage

For a model in which genotypes A_1_A_1_, A_1_A_2_ and A_2_A_2_ at a locus have fitness 1 − *s* and 1 − *t*, equations (6) and (7) can be replaced with:

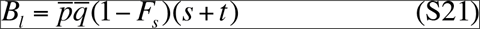

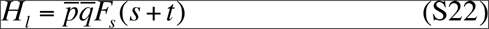

These expressions shows that the general principles established for mutation-selection balance apply to heterozygote advantage; if the effects of drift are moderate, so that the product of the two mean allele frequencies does not approach zero, both *B*_*l*_ and *H*_*l*_ will be of order (*s* + *t*) for intermediate values of *F*_*s*_. For the infinite island model with random mating, linearization around the deterministic equilibrium, *q*\* = *s*/(*s* + *t*), gives:

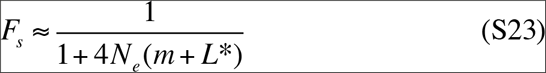

where *L*\* is the deterministic equilibrium genetic load with random mating, *st*/(*s* + *t*), which is also equal to the equilibrium value of *B*.

If *L*\* ≫ 1/(4*N*_*e*_) and *m, H*_*l*_ is approximately 1/(4*N*_*e*_), which is the same as the extra genetic load caused by drift in this case (Robertson, 1970), and is independent of the strength of selection and extent of population subdivision. With *L*\* ≪ 1/(4*N*_*e*_) and *m*, *H*_*l*_ is approximately *L*\**F*_*ST*_, and is thus proportional to both the extent of subdivision and the strength of selection.

---

**Table 1.**
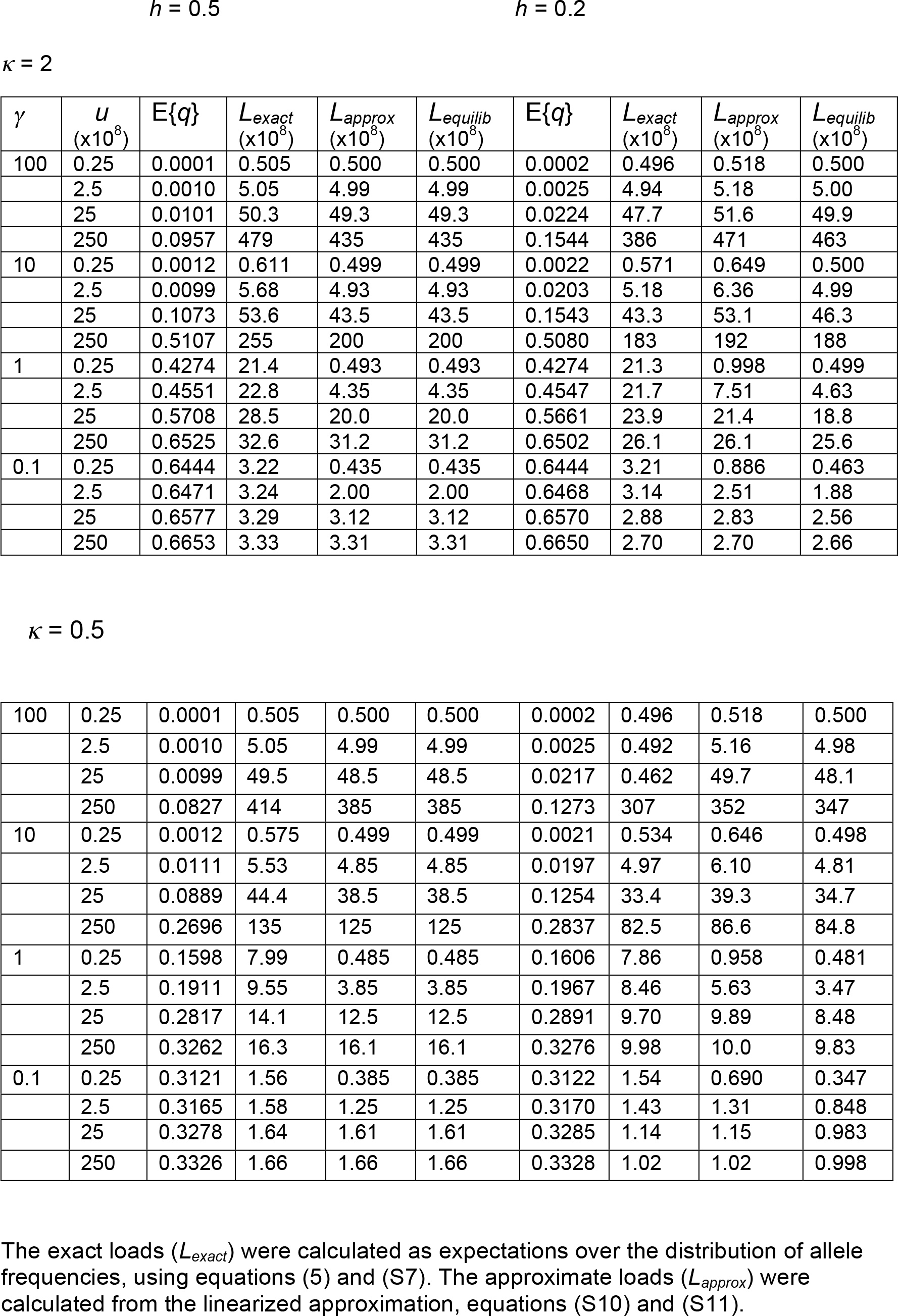
Genetic loads in a randomly mating population with drift, mutation and selection (*N*_*e*_ = 10^6^)

| $\gamma$ | $u$<br>($\times 10^8$) | $E\{q\}$ | $L_{exact}$<br>($\times 10^8$) | $L_{approx}$<br>($\times 10^8$) | $L_{equilib}$<br>($\times 10^8$) | $E\{q\}$ | $L_{exact}$<br>($\times 10^8$) | $L_{approx}$<br>($\times 10^8$) | $L_{equilib}$<br>($\times 10^8$) |
| --- | --- | --- | --- | --- | --- | --- | --- | --- | --- |
| 100 | 0.25 | 0.0001 | 0.505 | 0.500 | 0.500 | 0.0002 | 0.496 | 0.518 | 0.500 |
|  | 2.5 | 0.0010 | 5.05 | 4.99 | 4.99 | 0.0025 | 4.94 | 5.18 | 5.00 |
|  | 25 | 0.0101 | 50.3 | 49.3 | 49.3 | 0.0224 | 47.7 | 51.6 | 49.9 |
|  | 250 | 0.0957 | 479 | 435 | 435 | 0.1544 | 386 | 471 | 463 |
| 10 | 0.25 | 0.0012 | 0.611 | 0.499 | 0.499 | 0.0022 | 0.571 | 0.649 | 0.500 |
|  | 2.5 | 0.0099 | 5.68 | 4.93 | 4.93 | 0.0203 | 5.18 | 6.36 | 4.99 |
|  | 25 | 0.1073 | 53.6 | 43.5 | 43.5 | 0.1543 | 43.3 | 53.1 | 46.3 |
|  | 250 | 0.5107 | 255 | 200 | 200 | 0.5080 | 183 | 192 | 188 |
| 1 | 0.25 | 0.4274 | 21.4 | 0.493 | 0.493 | 0.4274 | 21.3 | 0.998 | 0.499 |
|  | 2.5 | 0.4551 | 22.8 | 4.35 | 4.35 | 0.4547 | 21.7 | 7.51 | 4.63 |
|  | 25 | 0.5708 | 28.5 | 20.0 | 20.0 | 0.5661 | 23.9 | 21.4 | 18.8 |
|  | 250 | 0.6525 | 32.6 | 31.2 | 31.2 | 0.6502 | 26.1 | 26.1 | 25.6 |
| 0.1 | 0.25 | 0.6444 | 3.22 | 0.435 | 0.435 | 0.6444 | 3.21 | 0.886 | 0.463 |
|  | 2.5 | 0.6471 | 3.24 | 2.00 | 2.00 | 0.6468 | 3.14 | 2.51 | 1.88 |
|  | 25 | 0.6577 | 3.29 | 3.12 | 3.12 | 0.6570 | 2.88 | 2.83 | 2.56 |
|  | 250 | 0.6653 | 3.33 | 3.31 | 3.31 | 0.6650 | 2.70 | 2.70 | 2.66 |
| $$\kappa = 0.5$$ |  |  |  |  |  |  |  |  |  |
| 100 | 0.25 | 0.0001 | 0.505 | 0.500 | 0.500 | 0.0002 | 0.496 | 0.518 | 0.500 |
|  | 2.5 | 0.0010 | 5.05 | 4.99 | 4.99 | 0.0025 | 0.492 | 5.16 | 4.98 |
|  | 25 | 0.0099 | 49.5 | 48.5 | 48.5 | 0.0217 | 0.462 | 49.7 | 48.1 |
|  | 250 | 0.0827 | 414 | 385 | 385 | 0.1273 | 307 | 352 | 347 |
| 10 | 0.25 | 0.0012 | 0.575 | 0.499 | 0.499 | 0.0021 | 0.534 | 0.646 | 0.498 |
|  | 2.5 | 0.0111 | 5.53 | 4.85 | 4.85 | 0.0197 | 4.97 | 6.10 | 4.81 |
|  | 25 | 0.0889 | 44.4 | 38.5 | 38.5 | 0.1254 | 33.4 | 39.3 | 34.7 |
|  | 250 | 0.2696 | 135 | 125 | 125 | 0.2837 | 82.5 | 86.6 | 84.8 |
| 1 | 0.25 | 0.1598 | 7.99 | 0.485 | 0.485 | 0.1606 | 7.86 | 0.958 | 0.481 |
|  | 2.5 | 0.1911 | 9.55 | 3.85 | 3.85 | 0.1967 | 8.46 | 5.63 | 3.47 |
|  | 25 | 0.2817 | 14.1 | 12.5 | 12.5 | 0.2891 | 9.70 | 9.89 | 8.48 |
|  | 250 | 0.3262 | 16.3 | 16.1 | 16.1 | 0.3276 | 9.98 | 10.0 | 9.83 |
| 0.1 | 0.25 | 0.3121 | 1.56 | 0.385 | 0.385 | 0.3122 | 1.54 | 0.690 | 0.347 |
|  | 2.5 | 0.3165 | 1.58 | 1.25 | 1.25 | 0.3170 | 1.43 | 1.31 | 0.848 |
|  | 25 | 0.3278 | 1.64 | 1.61 | 1.61 | 0.3285 | 1.14 | 1.15 | 0.983 |
|  | 250 | 0.3326 | 1.66 | 1.66 | 1.66 | 0.3328 | 1.02 | 1.02 | 0.998 |

The exact loads ( $L_{exact}$ ) were calculated as expectations over the distribution of allele frequencies, using equations (5) and (S7). The approximate loads ( $L_{approx}$ ) were calculated from the linearized approximation, equations (S10) and (S11).

**Table S2.** Estimates of *B* and *H* for natural populations of flowering plants.

| Species | Details | $B$ | $H$ | Reference |
| --- | --- | --- | --- | --- |
| *Phlox drummondii* | Self-incompatible; net fitness in field<sup>1</sup> | 0.31<br>(0.07) | 0.23<br>(0.09) | Levin & Bulinska-Radomska (1988) |
| *Scabiosa columbaria* | Outcrossing; net fitness in greenhouse<sup>2</sup> | 1.83<br>(0.84) | 0.76<br>(0.35) | Van Treuren et al. (1993) |
| *Salvia pratensis* | Outcrossing; survival x size in field<sup>3</sup> | 1.39<br>(0.20) | 0.32<br>(0.16) | Ouborg & Van Treuren (1994) |
| *Lychnis flos-cuculi* | Outcrossing; net fitness in common garden | 1.65<br>(0.19) | 0.18<br>(0.09) | Hauser & Loeschke (1994) |
| *Chamaecrista fasciculata* | Outcrossing; net fitness in field; Maryland 1995 data<sup>4</sup> | – | 0.18<br>(0.06) | Fenster & Galloway (2000) |
|  | Kansas 1995 data<sup>4</sup> | – | 0.18<br>(0.05) |  |
| *Gentianella germanica* | Self-compatible; net fitness in common garden; small populations | –0.33<br>(0.15) | 0.12<br>(0.07) | Paland & Schmid (2003) |
|  | Large populations | 0.12<br>(0.09) | 0.07<br>(0.07) |  |
| *Ranunculus reptans* | Self-incompatible; seed production in common garden<sup>5</sup> | – | 0.06<br>(0.05) | Willi & Fischer (2005) |
| *Leavenworthia alabamica* | Partially selfing populations; net fitness in greenhouse<sup>6</sup> | – | 0.39<br>(0.48) | Busch (2006) |
|  | Self-incompatible populations |  | 0.04<br>(0.18) |  |
| *Hypericum cumulicola* | Outcrossing; net fitness in greenhouse; small populations | –0.51<br>(0.31) | 1.20<br>(0.47) | Oakley & Winn (2012) |
|  | Large populations | 0.66<br>(0.45) | 0.07<br>(0.29) |  |
| *Arabidopsis lyrata* | Self-incompatible populations; net fitness in greenhouse | 1.02<br>(0.35) | 0.25<br>(0.18) | Willi (2013) |
|  | Partially selfing populations | 1.39<br>(0.79) | 0.96<br>(0.32) |  |
| *A. lyrata* | Self-incompatible populations; net fitness in greenhouse | 2.52<br>(0.60) | 0.47<br>(0.16) | Oakley et al. (2015) |
|  | Partially selfing populations | 1.67<br>(1.85) | 1.43<br>(0.75) |  |
| *Sabatia angularis* | Partially selfing; net fitness in common garden | 3.32<br>(0.82) | 0.74<br>(0.25) | Spigler et al. (2017) |

Standard errors of the means are in parentheses
<sup>1</sup> Mean and standard error of $B$ and $H$ obtained from values for 4 and 6 different crosses respectively.
<sup>2</sup> Mean and standard error of $H$ obtained from values for 6 different crosses.
<sup>3</sup> Mean and standard error of $B$ and $H$ obtained from values for 4 different crosses
<sup>4</sup> Mean and standard error of $B$ and $H$ obtained from values for 6 different crosses. The 1996 data for the Kansas field site had much smaller $H$ values, whereas the Maryland site had similar values to those in 1995.
<sup>5</sup> Mean and standard error obtained from values for 13 different crosses.
<sup>6</sup> Mean and standard error obtained from values for selfing and outcrossing populations for 3 and 4 different crosses, respectively.

For the other studies, the means were obtained by log-transforming the values of mean provided in tables or figures, and the standard errors were estimated from the ratios of the untransformed standard errors to the untransformed means.

**Figure S1.**
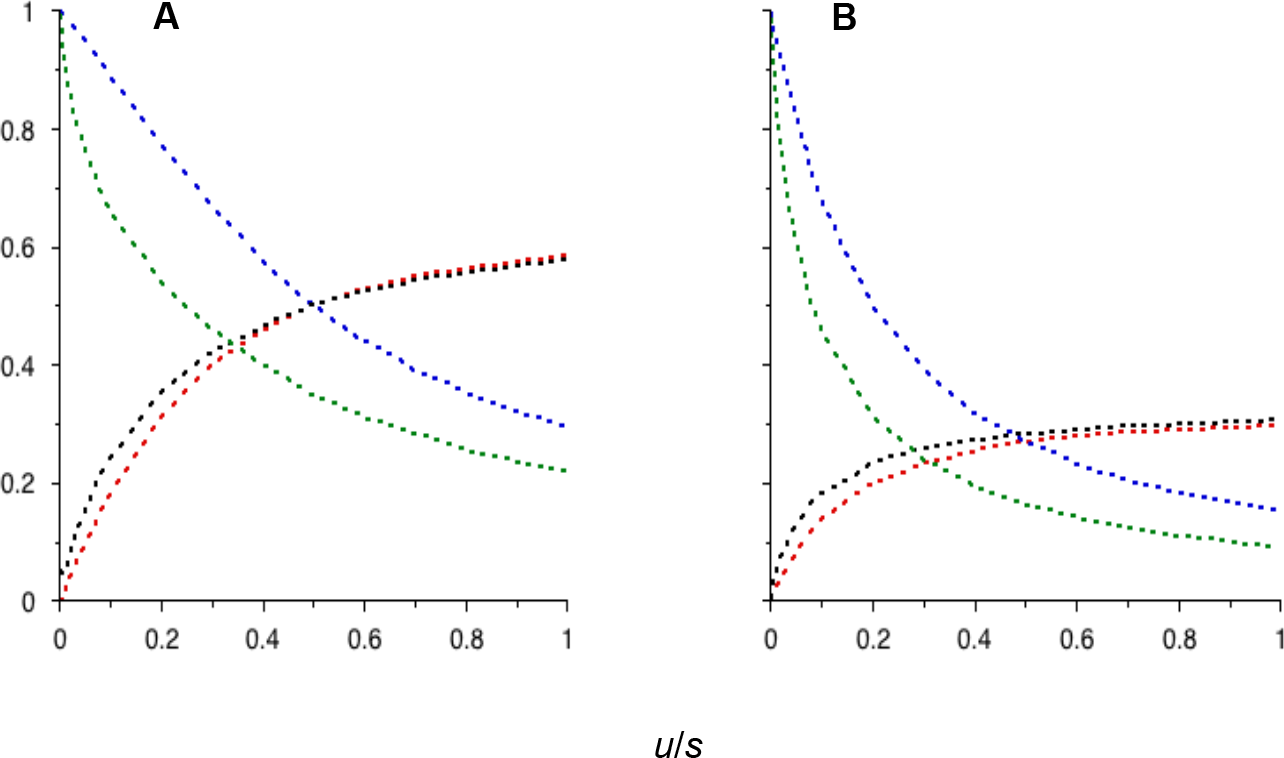
Plots of the deterministic equilibrium genetic load (green and blue curves, for *h* = 0.2 and *h* = 0.5, respectively), and the equilibrium frequency of the deleterious mutant allele (black and red curves, for *h* = 0.2 and *h* = 0.5, respectively), against the ratio of the mutation rate to the selection coefficient (*u*/*s*). Panel A is for the case of mutational bias in favour of the deleterious allele (*κ* = 2.0), and Panel B is for the case of mutational bias against it (*κ* = 0.5). The selection coefficient *s* against mutant homozygotes was 10^−4^. The genetic load was measured relative to 2*u*, the value expected for rare mutations with sufficiently strong heterozygous fitness effects that back mutations can be ignored.

**Figure S2.**
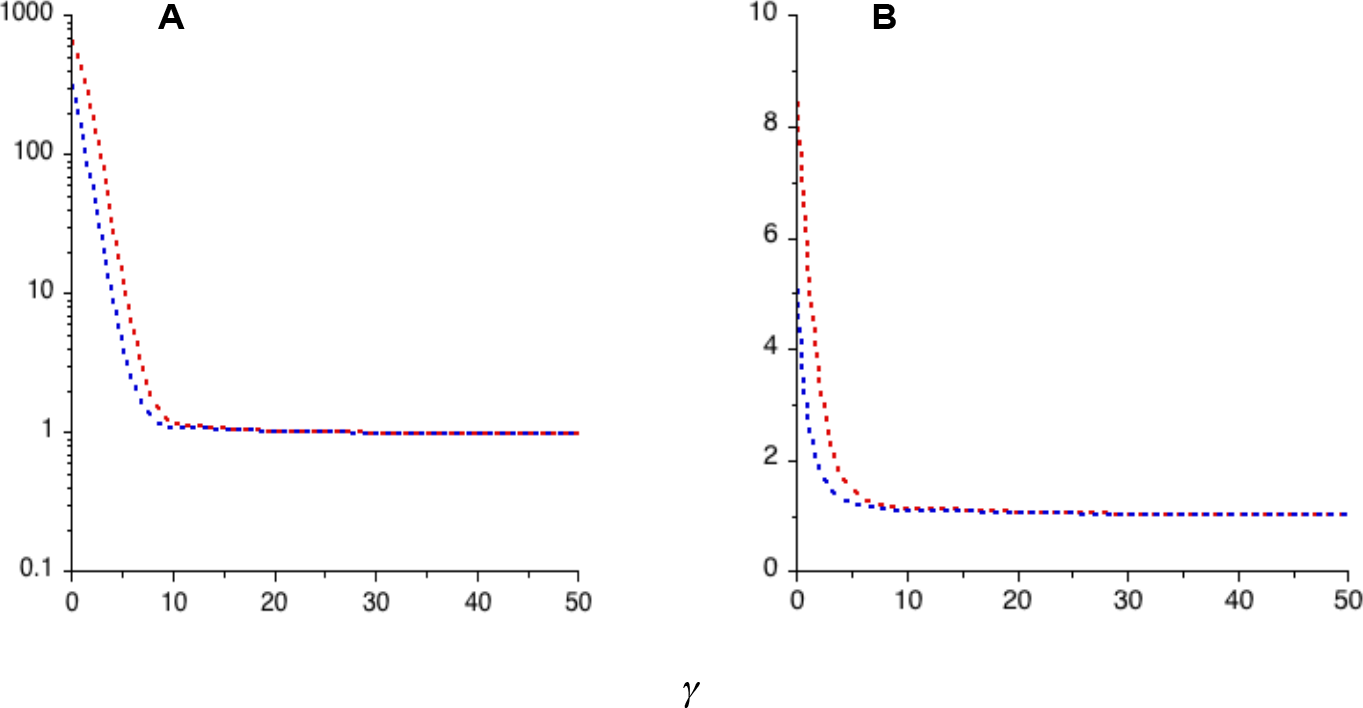
Plots of the mean genetic load at mutation-selection-drift equilibrium against *γ* = 2*N*_*e*_*s*, the scaled selection coefficient, which was altered by changing the population size while keeping *s* constant at 5 × 10^−6^. For the smallest value of *γ*, *N*_*e*_ = 10^4^. The load was measured relative to the exact deterministic equilibrium value, obtained from equations (3) and (4). The dominance coefficient, *h*, was 0.2. The blue and red curves are for *κ* = 0.2 and 0.5, respectively. Panel A is for a forward mutation rate of *u* = 2.5 × 10^−9^ and panel B is for *u* = 2.5 × 10^−7^.

